# NuMorph: tools for cellular phenotyping in tissue cleared whole brain images

**DOI:** 10.1101/2020.09.11.293399

**Authors:** Oleh Krupa, Giulia Fragola, Ellie Hadden-Ford, Jessica T. Mory, Tianyi Liu, Zachary Humphrey, Benjamin W. Rees, Ashok Krishnamurthy, William D. Snider, Mark J. Zylka, Guorong Wu, Lei Xing, Jason L. Stein

## Abstract

Tissue clearing methods allow every cell in the mouse brain to be imaged without physical sectioning. However, the computational tools currently available for cell quantification in cleared tissue images have been limited to counting sparse cell populations in stereotypical mice. Here we introduce NuMorph, a group of image analysis tools to quantify all nuclei and nuclear markers within the mouse cortex after tissue clearing and imaging by a conventional light-sheet microscope. We applied NuMorph to investigate two distinct mouse models: a *Topoisomerase 1* (*Top1*) conditional knockout model with severe neurodegenerative deficits and a *Neurofibromin 1* (*Nf1*) conditional knockout model with a more subtle brain overgrowth phenotype. In each case, we identified differential effects of gene deletion on individual cell-type counts and distribution across cortical regions that manifest as alterations of gross brain morphology. These results underline the value of 3D whole brain imaging approaches and the tools are widely applicable for studying 3D structural deficits of the brain at cellular resolution in animal models of neuropsychiatric disorders.

## Introduction

The mammalian cortex is composed of a diverse assembly of cell-types organized into complex networks, which function together to enable complex behaviors (Harris et al., 2019; Tasic et al., 2018; Zeisel et al., 2015). Disruption of cortical cytoarchitecture, either by genetic or environmental perturbation, can lead to altered brain function and create risk for neuropsychiatric disorders (Shin Yim et al., 2017; Stoner et al., 2014). A common approach for studying the mechanisms by which genetic variation increases risk for neuropsychiatric disorders is through the use of genetically modified animal models in a WT/KO experimental design. In order to observe the causal effects of disorder relevant genes on structure-function relationships, genetic tools can be applied to activate or silence genes in specific cell-types (Tsien et al., 1996) followed by imaging cellular organization through fluorescence microscopy. One critical goal in such experiments is to determine if the number of cells of a given type are altered by these genetic risk factors throughout different brain structures. However, a common limitation in imaging experiments done at cellular resolution is that they are restricted to anatomical regions of interest by physical sectioning which prevents the detection of region-specific effects. This becomes a particular issue for the cortex, one of the largest structures in the brain (Wang et al., 2020), where heterogeneity between cortical areas is often unmeasured by standard methods.

In order to image the entire brain without physical sectioning, tissue clearing methods render biological specimens transparent while preserving their 3 dimensional structure. Cleared tissues can then be rapidly imaged using light-sheet microscopy as plane illumination improves acquisition rates by 2-3 orders of magnitude compared to point scanning systems while also limiting the effects of photobleaching (Richardson and Lichtman, 2015; Ueda et al., 2020). Great strides have been made in the development of clearing protocols that are compatible with immunolabeling and the design of complementary sophisticated imaging systems (Matsumoto et al., 2019; Murray et al., 2015; Park et al., 2018; Susaki et al., 2020). Yet challenges still remain in expanding the accessibility of these technologies to research labs for quantitative analysis at cellular resolution.

For example, many of the current imaging protocols for whole brain profiling require custom light sheet systems to image tissues at cellular resolution (Fei et al., 2019; Matsumoto et al., 2019; Pende et al., 2018; Tomer et al., 2014; Voigt et al., 2019). These systems are therefore inaccessible to those lacking the expertise or resources required to assemble the necessary microscope components. Expanding tissues during the clearing process is a potential workaround that can increase the effective spatial resolution allowing for interrogation of subcellular structures without the need for custom imaging solutions (Chen et al., 2015; Gao et al., 2019; Ku et al., 2016; Murakami et al., 2018). However, expanded tissues can fall outside of the working distance of conventional microscope objectives, require prolonged imaging times, and significantly larger data storage resources. Therefore, computational tools designed for conventional light sheet microscope users are needed to compare cell counts in a WT/KO design.

With over 100 million cells in a mouse brain and image sizes of tissue cleared brain approaching terabytes, advanced image analysis tools are needed to achieve accurate cell quantification. Current segmentation methods for tissue cleared brain images apply a threshold for nuclear staining intensity and filter objects with a predefined shape, size, and/or density (Renier et al., 2016; Matsumoto et al., 2019; Fürth et al., 2018). However, variations in cell size, image contrast, and labeling intensity can all lead to inaccurate counts. In addition, whole brain images are typically registered to a standard reference, such as the Allen Reference Atlas (ARA), to assign cell locations to their corresponding structural annotations. Thus far, image registration has been performed mostly on stereotypical mice and has not been designed for mouse models with significant changes in gross morphology. With these limitations, the computational tools currently available have not been fully adopted for studying cellular organization in mouse models.

To address these issues, we developed a group of image analysis tools called NuMorph (Nuclear-Based Morphometry) (available here: https://bitbucket.org/steinlabunc/numorph/) for end-to-end processing to perform cell-type quantification within the mouse cortex after tissue clearing and imaging by a conventional light-sheet microscope. To demonstrate the effectiveness of the tool, we first applied and evaluated NuMorph to quantify structural changes in a mouse model with large differences in cortical structure, a topoisomerase I (*Top1*) conditional knockout (*Top1* cKO) mouse model that exhibits clear reductions in both cortical size and specific cell types (Fragola et al., 2020). We then apply NuMorph to investigate a neurofibromin I (*Nf1*) conditional knockout (*Nf1* cKO) model, a gene harboring mutations in individuals with Neurofibromitosis type I. This disorder often results in cognitive impairment, attention-deficit/hyperactivity disorder (ADHD), and autism spectrum disorder (ASD) (Gutmann et al., 2017). Our results reveal unique genetically influenced cell-type and structural changes in each mouse model, demonstrate the broad applicability of our analysis tools for studying both severe and subtle brain structure phenotypes in combination with tissue clearing methods, and present an alternative to 2D stereology for cellular quantification that does not rely on representative sampling.

## Methods

### Animals

*Top1* conditional knockout mice (*Top1*^*fl/fl*^*;Neurod6*-Cre, *Top1* cKO) were bred by crossing *Top1*^*fl/fl*^ mice (Mabb et al., 2016) with the *Neurod6*-Cre mouse line (Jackson Laboratory) (Goebbels et al., 2006) as described previously (Fragola et al., 2020). Cre-negative mice (*Top1*^*fl/fl*^) were used as controls (WT). *Nf1* conditional knockout mice (*Nf1*^*fl/fl*^) were purchased from (Jackson Laboratory, Stock No: 017639) (Zhu et al., 2001). *Emx1-Cre* mice were originally generated by Gorski et al. (Gorski et al., 2002) and previously validated and maintained in the lab (Xing et al., 2016). Homozygous *Nf1* cKO mice (*Nf1*^*fl/fl*^*;Emx1-Cre*) and heterozygous *Nf1* cKO mice (*Nf1*^*fl/+*^*;Emx1-Cre)* were generated by breeding *Nf1*^*fl/+*^;*Emx1-Cre* mice with *Nf1*^*fl/fl*^ *or Nf1*^*fl/+*^ mice. Cre-negative (*Nf1*^*fl/fl*^, *Nf1*^*fl/+*^, *Nf1*^*+/+*^) and *Nf1*^*+/+*^*;Emx1-Cre* mice were used as littermate controls. All animal procedures were approved by the University of North Carolina at Chapel Hill Institutional Animal Care and Use Committee. Mice were maintained on a 12-hr dark/light cycle and housed at temperatures of 18-23°C, 40-60% humidity, and *ad libitum* food and water. Genomic DNA extracted from tail or ear samples was utilized for genotyping by PCR. Primers for gene amplification are as follows (listed 5’-3’): *Top1-*F: GAGTTTCAGGACAGCCAGGA, *Top1*-R: GGACCGGGAAAAGTCTAAGC; Cre-F (*Neurod6*-Cre): GATGGACATGTTCAGGGATCGCC, Cre-R (*Neurod6*-Cre): CTCCCATCAGTACGTGAGAT, *Nf1*-F: ACATGGAGGAGTCAGGATAGT, *Nf1*-R: GTTAAGAGCATCTGCTGCTCT, Cre-F (*Emx1-Cre*): GAACGCACTGATTTCGACCA and Cre-R (*Emx1*-Cre): GATCATCAGCTACACCAGAG. Male P15 *Top1* cKO and WT littermate controls were used for tissue clearing in the *Top1* study. Male P14 *Nf1*^*fl/fl*^*;Emx1-Cre, Nf1*^*fl/+*^*;Emx1-Cre* and *Ctrl* littermates were used for tissue clearing in the *Nf1* study.

### Tissue Clearing & Immunolabeling

Tissue clearing was performed on 4 WT and 4 *Top1* cKO for the *Top1* study and 6 *Ctrl*, 6 *Nf1*^*fl/+*^;*Emx1-Cre*, and 6 *Nf1*^*fl/fl*^;*Emx1-Cre* for the NF1 study according to the iDISCO+ protocol (Renier et al., 2016). Genotyped samples were processed concurrently in littermate pairs/triplicates. Briefly, mice were fixed via transcardial perfusion using 4% paraformaldehyde and whole brain samples were dissected and cut along the midline. As the effects of *Top1* deletion on gross structure were bilateral upon visual inspection, only the left hemisphere was used in clearing experiments and analysis. Similarly, effects of *Nf1* deletion were found to be bilateral based on 2D stereological analysis and therefore only the right hemisphere was used in the *Nf1* study. Investigators were blinded to genotype for the *Nf1* study during tissue clearing and subsequent imaging and analysis. Large differences in overall brain size between *Top1* cKO and WT prevented blinding in this model. Samples were then washed in phosphate-buffered-saline (PBS), dehydrated in a graded series of methanol (Fisher, A412SK), pretreated with 66% dichloromethane (Sigma-Aldrich, 270997)/methanol and 5% H2O2 (Sigma-Aldrich, H1009)/methanol, followed by rehydration, permeabilization (20% dimethyl-sulfoxide, Fisher, BP2311; 1.6% Triton X100, Sigma-Aldrich, T8787; 23mg/mL Glycine, Sigma-Aldrich G7126), and blocking with 6% goat serum (Abcam, ab7481). Samples were then incubated with antibodies for Cux1 (Santa Cruz, sc-13024-Rb, 1:200) and Ctip2 (Abcam, ab18465-Rt, 1:500) for 5 days at 37°C in PTwH buffer (PBS; 0.5% Tween-20, Fisher, BP337; 10mg/L Heparin, Sigma-Aldrich, H3393). After 2 days of washing with PTwH, samples were then incubated with TO-PRO-3 (Thermo Fisher, T3605, 1:300), goat anti-rat Alexa Fluor 568 (Thermo Fisher, A11077, 1:200), and goat anti-rabbit Alexa Fluor 790 (Thermo Fisher, A11369, 1:50) for an additional 5 days at 37°C. Samples were then washed for 2 days with PTwH, dehydrated again using a graded methanol series, incubated in 66% dichloromethane/methanol for 3 hours, followed by a 30 minute incubation in 100% dichloromethane before storing in a dibenzyl ether solution (RI = 1.56, Sigma-Aldrich, 108014) at RT. Tissue clearing and antibody labeling required 21 days to complete.

### Light-Sheet Imaging

Imaging of cleared brain samples was performed using the Ultramicroscope II (LaVision Biotec) equipped with MVPLAPO 2X/0.5 NA objective (Olympus), sCMOS camera (Andor), and ImSpector control software. The zoom body was set to 2.5x magnification (yielding 1.21 μm/pixel) and a single light sheet was used with NA = ∼0.08 (9 μm thickness/ 4 μm z-step) as this allowed for better resolution of cell nuclei compared to using multiple light sheets. Dynamic horizontal focusing using the contrast enhanced setting in ImSpector was used to ensure axial resolution was maintained along the width of the image using the recommended number of steps depending on the laser wavelength. Samples were positioned sagittally with the cortex surface facing the single illuminating light-sheet (Figure S1D). This prevented excessive light scattering and shadowing from affecting the image quality in the cortical regions. Individual channels were acquired for tiled positions in a row-major order using 561nm (Ctip2), 647nm (ToPro), or 785nm (Cux1) laser lines. The 785nm channel was imaged first for the entire hemisphere. After refocusing the objective, the 561nm/647nm channels were then captured sequentially for each stack at a given tile position. Using these settings, mouse hemispheres were acquired using 3×3, 4×4, or 4×4 tiling schemes depending on hemisphere size with 5-15% overlap. Typical imaging times ranged from 10 to 15 hours for all 3 imaged channels. In the *Nf1* study, tissue autofluorescence was additionally imaged within a single tile using a 488nm laser at 0.8x magnification (3.86 x 3.86 x 4μm/voxel; 0.015 light sheet NA, no horizontal focusing) for all samples.

### Computing Resources

All data processing was performed locally on a Linux workstation running CentOS 7. The workstation was equipped with an Intel Xeon E5-2690 V4 2.6GHz 14-core processor, 8 x 64GB DDR4 2400 LRDIMM memory, 4 x EVGA GeForce GTX 1080 Ti 11GB GPU, and 2 x 4TB Samsung EVO 860 external SSDs. Hot swap bays were used to transfer data from the imaging computer to the analysis workstation.

### Image Preprocessing

Image preprocessing consists of all the necessary steps to prepare acquired raw images for image registration and cell quantification. All preprocessing steps were performed using custom written MATLAB R2020a scripts included in NuMorph and are described below.

### Intensity Adjustments

Two types of image intensity adjustments were performed on raw images prior to image stitching to increase accuracy of subsequent processing. First, uneven illumination along the *y* dimension (perpendicular to the light path) of each 2D image caused by the Gaussian shape of the light sheet was corrected using a MATLAB implementation of BaSiC, a tool for retrospective shading correction (Peng et al., 2017). We used 10% of all images, excluding tile positions around the cerebellum, to estimate a flatfield image for each channel. Each image was then divided by the flatfield prior to alignment and stitching to correct for uneven illumination. Second, differences in intensity distributions between image tile stacks, primarily as a result of photobleaching and light attenuation, were measured in the horizontal and vertical overlapping regions of adjacent tiles. To ensure bright features were of equal intensity between each stack, we measured the relative difference (*t* ^*adj*^) in the 95th percentile of pixel intensities in overlapping regions from 5% of all images. The measured image intensity *I*^*meas*^ at tile location (*x, y*)was then adjusted according to:

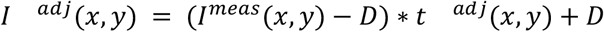

where *D* is the darkfield intensity (set as a constant value based on the 5th percentile of pixel intensities in all measured regions).

### Image Channel Alignment

As image channels are acquired one at a time, subtle drift in stage and sample positions during imaging may result in spatial misalignment between the reference nuclei channel and the remaining immunolabeled markers in a multichannel image. We tested two image registration approaches to ensure robust alignment across image channels. The first approach estimates 2D slice translations to align the immunolabeled channel images to the nuclear channel image. The axial (*z*) correspondence between the nuclei channel and every other channel within an image stack of an individual tile is first estimated using phase correlation at 20 evenly spaced positions within the stack. The correspondence along the axial direction with the highest image similarity (based on intensity correlation) determines the relative tile *z* displacement between channels (up to 50 μm in some cases). *xy* translations are then determined after multimodal image registration for each slice in the tile stack using MATLAB’s Image Processing toolbox. Outlier translations, defined as *x* or *y* translations greater than 3 scaled median absolute deviations within a local 10 image window in the stack, were corrected by linearly interpolating translations for adjacent images in the stack. In our data, outlier translations often occur in image slices without any sample present where the lack of image contents limits registration accuracy.

While a rigid 2D registration approach is sufficient for channel alignment when samples are securely mounted, sporadic movement of some samples during long imaging sessions can result in not only shifting translation but also rotational drift. In these cases, performing registration relying solely on translation will result in only part of the target image aligning correctly to the nuclei reference at a given *z* position with the remaining misaligned target features appearing in *z* positions immediately above and/or below (Figure S1B). To correct for these displacements, we applied a nonlinear 3D registration approach using the Elastix toolbox (Klein et al., 2010) between channels for each individual tile. Full image stacks were loaded and downsampled by a factor of 3 for the *x/y* dimensions to make the volume roughly isotropic and reduce computation time. Intensity histogram matching was then performed and a mask was identified for the nuclei reference channel using an intensity threshold that limits sampling positions in the background. Next, an initial 3D translational registration is performed on the entire image stack between the reference and the remaining channels. The stack is then subdivided into smaller chunks of 300 images and rigid registration is performed on each chunk to account for 3D rotation and achieve a more accurate initial alignment within local regions of the full stack. Finally, a nonlinear B-spline registration is performed on each chunk using an advanced Mattes mutual information metric to account for *xy* drift along the *z* axis and ensure precise alignment of image features. B-spline transformation grid points were set to be sparser along *xy* compared to *z* (800×800×8 voxels) as this setting well balances accurate alignment with computational cost while also preventing local warping of background intensities.

During image processing, the 2D rigid alignment approach was initially used to align each sample. Each tile was then visually inspected to ensure accurate alignment of all channels along the stack. For tiles where rigid alignment was inaccurate, the non-rigid alignment method was used to correct for misalignment.

### Iterative Image Stitching

A custom 2D iterative stitching procedure was used to assemble whole brain images at high resolution. First, an optimal pairwise *z* correspondence along the axial direction was determined for adjacent tile stacks by exhaustive image matching for the horizontally and vertically overlapped candidate regions. Specifically, a sample of 10 evenly spaced images were taken within a stack and registered to every *z* position within a 20 image window in the adjacent stack using phase correlation. The displacement in *z* with the highest count of peak correlations among the 10 images was presumed to represent the best *z* correspondence. The difference in correlation between the best and the 2^nd^ best *z* displacement was used as a weight for the strength of the correspondence, with a larger difference representing a stronger correspondence. This resulted in 4 matrices: pairwise horizontal and vertical *z* displacements and their corresponding weights. To determine the final *z* displacement for each tile, we implemented a minimum spanning tree (Kruskal, 1956) using displacements and their weights as vertices and edges, as previously implemented (Chalfoun et al., 2017).

An intensity threshold to measure the amount of non-background signal was determined by uniformly sampling 5% of all images and calculating the median intensity. The starting point for iterative stitching going up/down the stack was selected at a position near the middle of stack with sufficient non-background signal (set to 1 standard deviation above the darkfield intensity) present in all tiles. Translations in *xy* were calculated using phase correlation and further refined using the Scale Invariant Feature Transform (SIFT) algorithm (Lowe, 2004). The top left tile was set as the starting point for tile placement for each stitching iteration. This ensures stitched images would not be shifted relative to each other along the *z* axis. Tiles were blended using sigmoidal function to maintain high image contrast in overlapping regions. Spurious translations, defined as translations greater than 5 pixels in *x* or *y* from the previous iteration, in images that lacked image content were replaced by translation results from the previous iteration.

### Image Registration to ARA Using Point Correspondence

Volumetric image registration was performed using Elastix to measure the correspondence between the stitched TO-PRO-3 channel in the tissue cleared samples and the Nissl-stained Allen Reference Atlas (ARA) (Dong, 2008; Lein et al., 2007). The atlas and corresponding volume annotations from Common Coordinate Framework v3 were downloaded using the Allen Software Development Kit (SDK) (https://allensdk.readthedocs.io/) at 10 μm/voxel resolution. In each registration procedure, the ARA was downsampled to 25 μm/voxel resolution to perform registration and the resulting transformation parameters were rescaled and applied to the annotation volume at the native 10 μm/voxel resolution.

For registration without point guidance, an affine followed by B-spline transformation sequence was applied along 3 resolution levels to each sample using advanced mattes mutual information (MMI) as the sole metric to estimate spatial correspondence (as done previously in (Renier et al., 2016). This registration procedure allowed for direct mapping of ARA annotations to each registered sample and was applied to all WT hemispheres in the *Top1* study. Adding point guidance to WT samples resulted in similar registration accuracy but slightly higher variation in structure volumes between samples (Figure S3).

A modified version of the standard registration procedure without points was also used for mapping control and *Nf1* cKO hemispheres in the *Nf1* study as this knockout model exhibited a lower degree of morphological variation compared to the *Top1* cKO model. However, to further improve registration accuracy, we incorporated additional spectra from tissue autofluorescence that was mapped to the ARA average MRI template, in addition to the TO-PRO-3/Nissl mapping. The downsampled autofluorescence channel in each sample was initially pre-aligned to the TO-PRO-3 reference using rigid registration and the standard B-spline registration with the ARA Nissl/MRI templates proceeded while maximizing the joint mutual information correspondence between the channel pairs.

For points-guided registration in the *Top1* cKO model, we first manually placed 200 landmarks within both the ARA and our to-be-registered nuclei reference image, using the BigWarp plugin in Fiji (Bogovic et al., 2016). The majority of points were located within or around the cortex, as this was our region of interest and contained the largest deformations in the *Top1* cKO samples (Figure S4). The same set of reference point coordinates in the ARA were selected for each sample and used as input points in Elastix for affine and B-spline registration along 3 resolution levels. Estimates of spatial correspondence for points-guided registration was driven by a hybrid metric based on (1) minimizing the point distances between two images and (2) maximizing the voxel-wise image similarity between two images which is measured by mattes mutual information (MMI). For affine registration, voxel-wise similarity (based on MMI) was ignored and only points distance was used to estimate global translation, rotation, and scaling transformations. For B-spline registration, we gradually increased the influence of voxel-wise similarity in the hybrid metric during the registration sequence from coarse to fine resolution (1:0.2, 1:0.4, 1:0.6; MMI:Point Distance weight). The inverse of the final transformation parameters was then calculated using a displacement magnitude penalty cost function (Metz et al., 2011) and applied to the Allen Mouse Brain Common Coordinate Framework v3 annotation volume to assign anatomical labels for each voxel in the native sample space. While a more direct approach would be to register the ARA to the sample, we found that registering the sample to the ARA and calculating the inverse achieved slightly higher accuracy in *Top1* cKO brains (data not shown).

To evaluate registration accuracy, 3D masks of the entire isocortex were manually labeled for each sample in Imaris (Bitplane) using the 3 acquired channels as markers to delineate cortex boundaries. Some cortical subplate structures, such as the claustrum, were included in the final mask as these were difficult to distinguish from the isocortex. The DICE similarity score was then calculated between each mask and all cortical structures in the registered annotation volume (Figure 2B) as a metric of registration accuracy.

### Cortical Volume, Surface Area, and Thickness Measurements

Quantitative measurements for the volume, surface area, and thickness of the isocortex and 43 cortical areas defined in (Harris et al., 2019) or a lower level set of 17 cortical areas (ARA structure depth=6) were calculated based on registered annotation volumes. The voxel sums (at 10 um^3^/voxel) represent the total volume of each structure. To calculate volumetric displacement for each sample relative to the Allen atlas, the spatial Jacobian was measured for each set of transformation parameters, which ranges from -1 to 1, and represents voxel-wise local compression or expansion. Surface area for the isocortex was calculated based on MATLAB’s implementation of Crofton’s formula (Lehmann and Legland, 2012). The fraction of layer 1 boundary voxels over all boundary voxels was used to determine the area of only the outer cortical surface. This measurement was then further partitioned by the number of layer 1 boundary voxels for each individual structure. To calculate thickness, the center of mass for layer 1 and layer 6b were first calculated for each structure. Thickness was then measured based on the euclidean distance between 2 points within layer 1 and layer 6b that were nearest to the centers of mass. Average thickness of the full isocortex was weighted by the volume contribution of each structure.

### Nuclei Detection

Imaging data for training the 3D-Unet model was acquired from 3 separate imaging experiments of TO-PRO-3 labeled nuclei across 5 different regions from the cortex of 2 WT brains. Images were captured at 0.75×0.75×2.5 μm/voxel for training a high resolution model or 1.21×1.21×4 μm/voxel for training a low resolution model. A binary approximation of the nucleus volume was initially pre-traced using the cell detection component of the CUBIC-informatics pipeline (Matsumoto et al., 2019). Specifically, the thresholded Hessian determinant after Difference-of-Gassian filtering was used to create an initial 3D mask of all nuclei in the image. Full images were then divided into patches of 224×224×64 voxels and preprocessed using min/max normalization. The corresponding 3D mask for each nucleus was reduced to its 2D component at the middle z position. Each patch was then manually inspected and corrected for segmentation error or incorrect shapes using BrainSuite v17a (Shattuck and Leahy, 2002) by 1 rater (OK) to reduce person-to-person variability. The corrected 2D nuclei masks were then eroded by removing 40% of the outer edge pixels. Each patch was then subdivided into 4 smaller patches of 112×112×32 voxels, with 1 out of the 4 patches being withheld for the validation set. The full dataset (training + validation) contained 16 patches at 224×224×64 voxels for both the high (14,554 nuclei) and low resolution (53,993 nuclei) models. Nuclei at the edge of an image stack were also included in the training. Manually labeled data are available at https://braini.renci.org/ using the Download Image service.

A modified 3D-Unet architecture (Çiçek et al., 2016; Isensee et al., 2018) was used to identify the positions of cell nuclei in whole cortex images. We built upon and modified a previous Keras implementation of 3D-Unet for volumetric segmentation in MRI (https://github.com/ellisdg/3DUnetCNN) to detect binary masks of cell nuclei positions. As originally described (Isensee et al., 2018), the 3D-Unet architecture contains a series of context modules during the contracting path that encodes abstract representations of the input image, followed by a series of localization modules on the upscaling path to localize the features of interest (Figure S4A). We similarly used a model with 5 context modules, residual weights, and deep supervision in the localization modules. The network was trained using 32 base filters on image patches of size 112×112×32 voxels with a batch size of 2. Training presumed over ∼300 epochs using an Adam optimizer with a dropout rate of 0.4 and an initial learning rate 0.002 that was reduced by a factor of 2 for every 10 epochs without the loss improving. Additional image augmentations were implemented during the training to make the model more generalizable. These include random image permutations, image blurring and sharpening, the addition of random noise, and intensity variations along *x,y,z* dimensions in the image patch. Random scaling was removed as we found that this decreased model performance.

Nuclei detection accuracy was evaluated using an independent set of 5 images patches of TO-PRO-3-labeled nuclei where the full 3D volume of each nucleus was fully manually drawn with a unique index at 0.75×0.75×2.5 μm/voxel resolution (∼3,500 nuclei total). Each patch was sampled from a unique region within 1 WT cortex. Evaluation patches were initially delineated by 4 raters and further refined by 1 rater to reduce between-rater variability. We compared our 3D-Unet detection method with those used in 2 previously published pipelines for tissue cleared image analysis: ClearMap and CUBIC-informatics (Matsumoto et al., 2019; Renier et al., 2016). For ClearMap, we used voxel size and intensity thresholds after watersheding, as described in the published implementation. Parameters for cell size and intensity were scaled accordingly to achieve the most accurate average cell counting results possible for all the patches tested. Similarly, intensity normalization and Difference-of-Gaussian scaling parameters used in CUBIC-informatics were adjusted according to image resolution. Filtering by intensity and structureness was also performed as described in the previous work (Matsumoto et al., 2019).

In our evaluation of nuclei detection, precision is the proportion of nuclei correctly predicted out of all nuclei predictions in an image patch. Precision is therefore calculated by counting the number of cells with multiple predicted centroids in 1 manually labeled nucleus volume as well as false positives cells called in the image background divided by the total number of nuclei detected and subtracting this number from 1. Recall is the proportion of all nuclei instances that were predicted. Recall was therefore calculated by counting the number of manually labeled cell volumes that lacked any predicted cell centroids divided by the total number of cells. The majority of false negative cases were due to touching nuclei. Nuclei whose centroid were within 3 voxels of the image border were excluded from the evaluation.

Whole brain TO-PRO-3 images were divided into chunks of 112×112×32 voxels to be fed into the trained 3D-Unet model for prediction of cell centroids. An overlap of 16×16×8 voxels was used between adjacent chunks to minimize errors from nuclei at chunk edges. Centroid positions falling in a region less than half the overlap range (i.e. <8 pixels from *xy* border or <4 pixels from *z* border) were assumed to be counted in the adjacent overlapping chunk and were removed. Additionally, a nearest neighbor search using kd-trees (Bentley, 1975) was performed to remove duplicate centroids within 1.5 voxels of each other, ensuring centroids in overlapping regions were not counted multiple times. Increasing overlap did not significantly affect the final cell counting results (data not shown). Total computation time for detecting all cortical nuclei in 1 WT brain hemisphere was ∼2.5 hours using a single GPU.

### Cell-type Classification

To classify cell-types, we took a supervised approach by training a linear Support Vector Machine (SVM) classifier using MATLAB’s Statistics and Machine Learning Toolbox on a set of intensity, shape, and annotation features within a 2D patch surrounding each centroid. First, channel intensities were measured at centroid positions for each channel. Cells with intensities below the median for both Ctip2 and Cux1 were presumed negative for both markers and removed from model training and classification (∼25% of cells). In the remaining cells, we took a uniform, random sample of 1,000 cells from each brain image dataset and retained 2D patches (13×13 pixels) around centroid positions. Manual classification required >1 hour per dataset using a custom NuMorph function that allows fast navigation between cell patches. For each patch, we recorded several intensity measurements (max, mean, standard deviation, middle pixel, middle pixel/edge pixel) and applied Otsu thresholding to capture shape measurements (total filled area, inner filled area) in each channel. These were also combined with categorical annotations for cortical layer (L1, L23, L4, L5, L6a, L6b) and cortical area (Prefrontal, Lateral, Somatomotor, Visual, Medial, Auditory; defined in (Harris et al., 2019). Cells were then manually classified into 4 classes: (1) Ctip2-/Cux1-, (2) Ctip2+/Cux1-, (3) Ctip2-/Cux1+, (4) Outlier. The outlier class was annotated according to 4 additional subdivisions due to differences in intensity features: (1) Ctip2+/Cux1+, (2) Pial surface cell, (3) TO-PRO-3-/Ctip2-/Cux1-(4) Striatal cell (only present in *Top1* cKO from residual registration error near white matter boundary). The SVM model was then trained using all intensity, shape, and annotation features. Model accuracy was evaluated using 5-fold cross-validation and applied to the remaining cells for classification. Due to differences in labeling intensity between samples, we trained a new model for each sample instead of aggregating annotation data.

We compared supervised cell classification with an unsupervised approach based on modeling fluorescence intensities at centroids positions as Gaussian mixtures (GM) for Ctip2 and Cux1. After Z normalization, high intensity cells (Z > 5 and Z < -5) winsorized and outliers expressing both markers near the sample edge were removed. GM model fitting was then performed separately on normalized Ctip2 and Cux1 intensities using 2 or 3 components (whichever had higher accuracy by visual inspection) for 20 replicates using parameters initialized by k-means++ (David Arthur, 2007). Due to spatial variation in gene expression, we stratified GM model fitting to 6 general areas defined in (Harris et al., 2019) according to each cell’s structural annotation to further improve accuracy. We then calculated posterior probabilities of each cell being positive for either marker. Cells with a posterior probability greater than 0.5 of not being background were classified as positive. As the vast majority of neurons do not co-express Ctip2 and Cux1 (Molyneaux et al., 2007), we filtered Ctip2+/Cux1+ cells according to their layer annotation. Cells in L1-L4 with P(Cux1) > P(Ctip2) were classified as Cux1+ and cells in L5-L6b with P(Ctip2) > P(Cux1) were classified as Ctip2+. The remaining Ctip2+/Cux1+ cells were classified as outliers.

### Quantification, Statistical Analysis, and Visualization

Final cell-type counts were summed for each annotation in the cortex according to its structure tree hierarchy. In our analysis, we chose to compare either 43 cortical areas defined in (Harris et al., 2019) or a lower level grouping of 17 regions based on the ARA structure hierarchy. Statistics, including mean counts, standard deviation, fold change, raw p values, and false discovery rate (FDR) adjusted p values (Benjamini-Hochberg; FDR < 0.05), were calculated in MATLAB and exported for plotting using custom R scripts and customized slice visualization. Structure volumes were also used to calculate cell density statistics. Unless stated otherwise, descriptive statistics in the main text and error bars in figure plots represent mean ± standard deviation.

2D slice visualizations were created using a custom MATLAB program based on the allenAtlasBrowser in the SHARP-Track tool (Shamash et al., 2018). Structure annotations were downsampled along the anterior-posterior axis to reduce memory overhead for smoother performance and colored by volume, cell count, or cell density statistics. Additional visualizations for point clouds, surface volumes, and flattened isocortex plots were created using custom MATLAB scripts and are available in the NuMorph package. Additional animations were generated in Imaris (Bitplane) after importing cell centroid position as “spots” objects.

### Spatial Gene Expression Correlation

Fold change in cell counts between WT and *Top1* cKO were correlated with spatial gene expression based on *in situ* hybridization measurements from the Allen Mouse Brain Atlas (Lein et al., 2007). Expression grid data from sagittal and coronal sections were downloaded using the Allen SDK. Expression energy for each gene was first Z-scored across all brain structures and cortical regions were retained for analysis. Duplicate sections for the same gene were combined by taking the mean Z score for each structure across sections. We filtered out any gene that did not have expression data in all cortical structures and removed genes with Z scores less than 1 in all structures as these represent genes with consistently low cortical expression or with low congruence between duplicate sections. For the remaining genes, we applied a robust sigmoidal transformation as described in (Fulcher and Fornito, 2016) to account for the presence of outliers in ISH expression data. As certain cortical regions also have greater cell density and therefore greater total ISH energy, we conducted an additional Z score normalization across cortical regions to have the same average total gene expression.

To reduce known false positive associations from gene-gene coexpression (Fulcher et al., 2020), we ran comparisons to ensemble-based random null models generated using the Gene Category Enrichment Analysis toolbox (https://github.com/benfulcher/GeneCategoryEnrichmentAnalysis). Null distributions were generated for GO categories containing between 10 and 200 genes by 10,000 random samples to create a Gaussian distribution estimate of each GO null distribution. In total, we used null models for 4,186 GO categories based on expression of 10,945 genes across 38 cortical structures. Correlations between spatial gene expression and relative cell count differences were tested and corrected for multiple-hypothesis testing using a false discovery rate of 0.05.

Additional annotations for gene length comparisons were downloaded from Ensembl (Cunningham et al., 2019). The Spearman correlation between each gene’s expression and cell count or density differences across cortical regions was measured and binned by gene length based on the longest isoform for each gene. The mean and standard deviation of all correlation coefficients in each bin (<100kb or >100kb) was used to compare correlation coefficients between bins (Welch’s t-test). A list of differentially expressed genes in *Top1* cKO cortex as measured by scRNA-seq was acquired from (Fragola et al., 2020) for additional comparisons.

### Brain Section Preparation and Immunofluorescence Staining

For histological studies, mice were anesthetized and perfused transcardially with 4% paraformaldehyde (PFA) /1XPBS. Brains were dissected and postfixed in 4% PFA for 16 hours. Brains were embedded in 4% low-melting point agarose/1XPBS and sectioned using Leica VT1200 vibratome. Sections were stored in 1XPBS at 4°C.

For immunohistology studies, sections were rinsed in PBS and incubated in blocking solution (5% normal serum, 0.3% Triton X-100, 2% DMDO, 0.02% Sodium Azide,1XPBS) at room temperature. Primary antibodies were diluted in blocking solution and incubated overnight at room temperature. The following antibodies were utilized for immunofluorescence: rabbit anti-Cux1 (Santa Cruz, sc-13024; 1:500), rat anti-Ctip2 (Abcam, ab18465; 1:1000), rabbit anti-Satb2 (Abcam, ab51502; 1:1000), goat anti-GFAP (Abcam, ab53554; 1:1000) and rabbit anti-Olig2 (Millipore, AB9610; 1:2000). Brain sections were rinsed in 0.1% Triton-X 100/1XPBS (PBS/T) three times and incubated with secondary antibodies in blocking solution for 3 hours at room temperature. Secondary antibodies utilized include donkey anti-rabbit IgG Alexa Fluor 568 (Thermo Fisher, A10042; 1:1000), goat anti-rat IgG Alexa Fluor 488 (Thermo Fisher, A11006; 1:1000) and donkey anti-goat IgG Alexa 488 (Thermo Fisher, A11055; 1:1000). Sections were then stained with DAPI (1:1000 in PBS/T) and rinsed with PBS/T three times for 20 minutes each. Sections were mounted onto Fisherbrand Superfrost/Plus slides with antifading Polyvinyl alcohol mounting medium with DABCO (Sigma-Aldrich, 10981). Images were collected with a Zeiss LSM 780 laser scanning confocal microscope.

### EdU Labeling and Detection

To permanently label newly generated cortical neurons, pregnant dams at E16.5 were single-dosed with 5–ethynyl–2′–deoxyuridine (EdU, Cayman Chem) in 1X PBS at 30mg/kg body weight via intraperitoneal injection. Pups were perfused at P14 and sectioned as described above. EdU detection was conducted upon the completion of immunofluorescence labeling. After washing in PBS for 10 minutes, brains sections were incubated with Alexa 647-conjugated Azide (Biotium) at 1.6µM, in solution containing 0.1M Tris-HCl (pH 8.8), 0.43 X PBS, 4 mM CuSO_4_, and 0.1M Ascorbate, for 20 minutes. Brains sections were washed with PBS/T three times at 20 minutes each, stained with DAPI and mounted for confocal imaging.

### Confocal Image Analysis and Quantification

Confocal images of barrel fields from 2D slices were collected for cortical thickness and cell number analysis. Images were acquired from anatomically matched coronal sections along the rostro-caudal axis. The distribution of cortical upper layer (layer 2-4) marker, Cux1, was utilized to determine the boundaries separating layer 1, upper layers and lower layers (layer 5-6). The thickness of an individual layer was measured along the middle segment of selected regions of interests (ROIs). For cell number assessment, images were processed using ImageJ (https://imagej.net/ImageJ). Briefly, images were auto-thresholded at default setting with manual adjustment to eliminate unfocused signals and binary images were watershedded. Numbers of Cux1, Ctip2 and Olig2 expressing cells were automatically determined using the Analyze Particles function with a cut off size at 17.5µm^2^. To determine EdU-labeled cortical neurons, EdU and Satb2 co-labeled cells were extracted and cell numbers were automatically determined using the Analyze Particles function with a cut off size at 35.0µm^2^. GFAP expressing cells in the cortical plate were counted manually in Photoshop after thresholding using ImageJ. Representative images were cropped and adjusted for brightness and contrast in Photoshop for presentation. Mice from a minimum of three litters were analyzed for each experiment. 2 to 3 ROIs were analyzed and results were averaged from each animal. For all experiments, n represents the number of animals. One-way ANOVA analyses with *post hoc* Tukey’s tests were performed in GraphPad Prism 9.

### Data and Code Availability

NuMorph source code is available at https://bitbucket.org/steinlabunc/numorph/. Manually labeled annotations for 3D-Unet training and raw light-sheet images are available at https://braini.renci.org/ through the “Download Image” service.

## Results

### iDISCO+ Reveals Neuronal Cell-type Deficits in the *Top1* cKO Cortex

A previous study demonstrated that deletion of *Top1* in postmitotic excitatory neurons within the cortex and hippocampus results in massive neurodegeneration in these structures by postnatal day 15 (P15) (Fragola et al., 2020). Interestingly, while all cortical layers were affected by *Top1* deletion, the lower cortical layers (Layers 5-6) showed a noticeably greater reduction in thickness and cell count compared to the upper cortical layers (Layers 2-4) (Fragola et al., 2020). These observations however were limited to the somatosensory cortex, which itself is a large structure that can be further decomposed into multiple functional regions. To evaluate the effects of *Top1* deletion on excitatory neuron cell-types throughout all cortical structures, we performed iDISCO+ (Renier et al., 2016) to clear and image the *Top1* cKO (*Neurod6*^*Cre/+*^::*Top1*^*fl/fl*^) mouse. We chose to use iDISCO+ among other tissue clearing techniques due to its demonstrated compatibility with antibody labeling, minimal tissue expansion or shrinkage, and simplified protocol (Renier et al., 2016). To go beyond qualitative evaluation, we proceeded to develop cell detection and image registration tools that could accurately quantify the number of upper layer and lower layer neurons in each cortical region in *Top1* cKO mice (Figure 1A).

**Figure 1.**
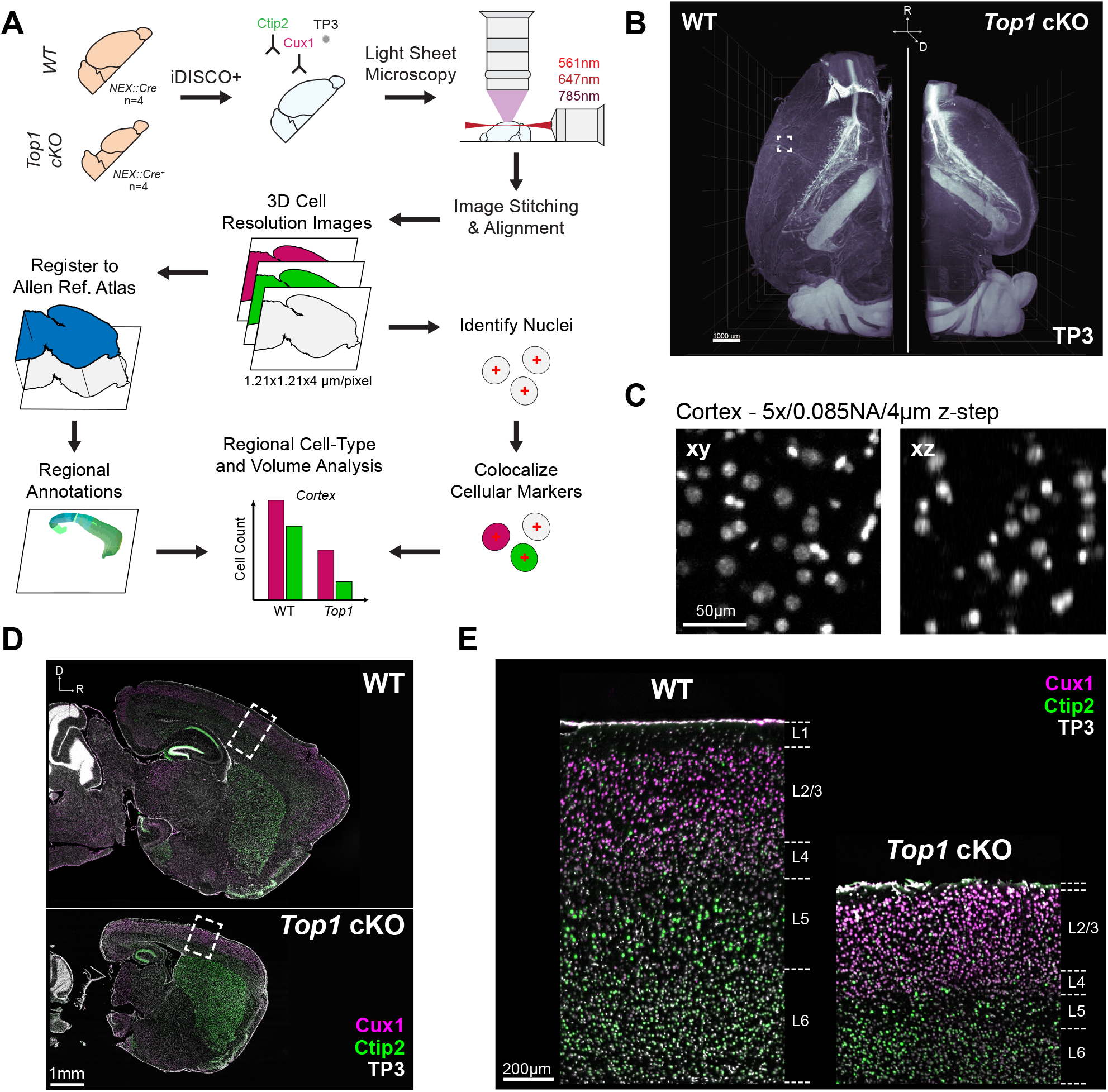
Cellular Resolution Analysis of Brain Structure Phenotypes for Tissue Cleared 3D Brain Images. A. Overview of tissue processing, imaging, and image analysis procedures. B. 3D rendering of cell nuclei in WT and *Top1* cKO samples. C. Example of TO-PRO-3 (TP3) labeled nuclei within WT cortex captured at sufficient lateral (*xy*) and axial (*xz*) resolution for cell quantification. D. Optical sagittal sections of TO-PRO-3 nuclear staining and immunolabeling for cell-type specific markers Ctip2 (lower layer neuron) and Cux1 (upper layer neuron) in WT and *Top1* cKO samples. E. Zoomed in images of boxed cortical areas in D demonstrating channel alignment and showing the expected localization of upper and lower layer markers.

**Figure 2.**
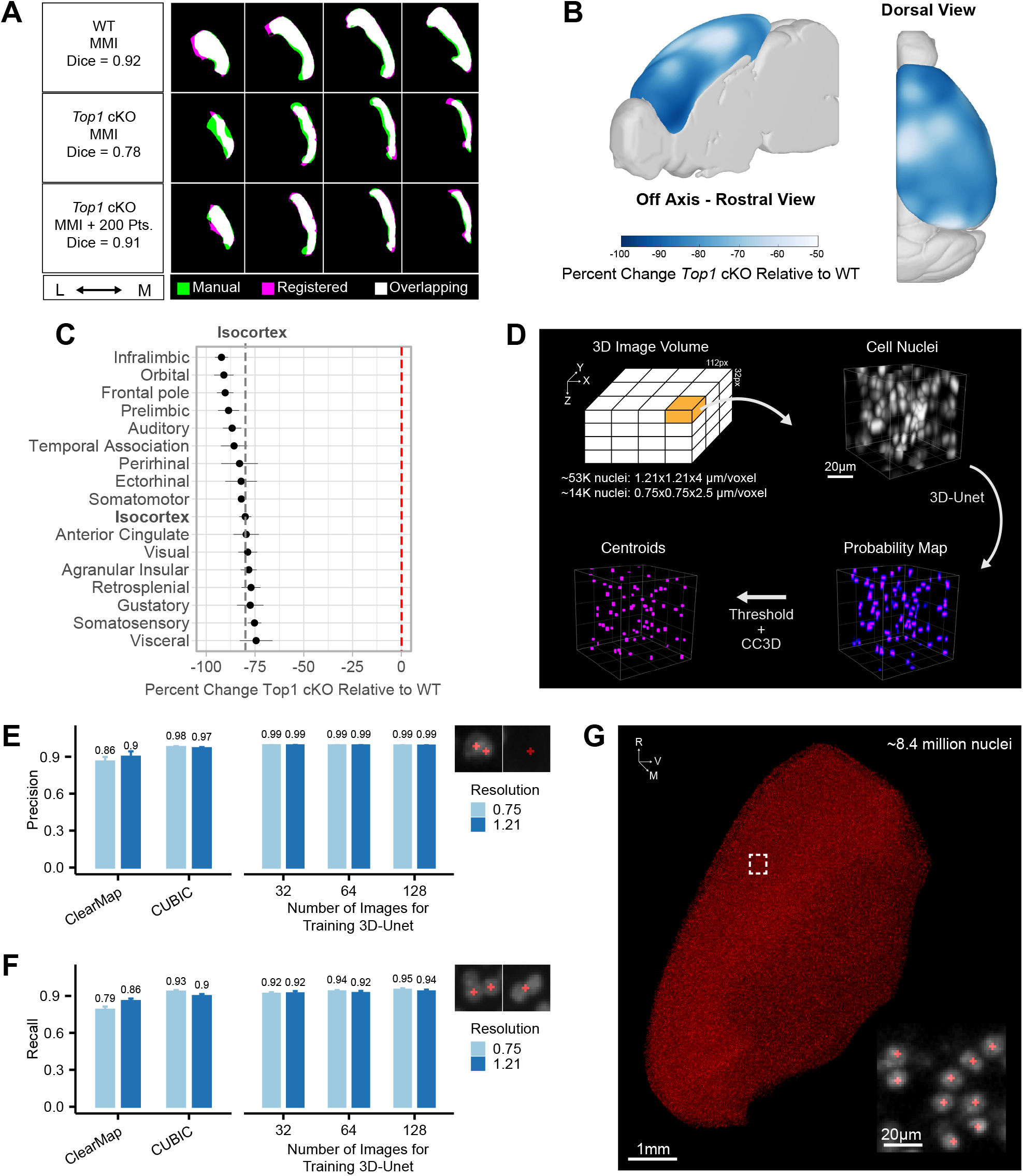
NuMorph Integrates Point Correspondence to Register Difficult Structures and 3D-Unet for Accurate Detection of Cortical Nuclei. A. Cortical masks from registered WT and *Top1* cKO brain images (Magenta) compared with manual labelled traces (Green). Mattes mutual information (MMI) was used as the primary registration metric with additional point correspondence to guide registration in the *Top1* cKO case. B. Voxel-wise differences in cortical volumes between *Top1* cKO and WT samples. C. Percent change in cortical region volumes in *Top1* cKO samples compared to WT. Dashed line indicates average change across the entire cortex. Data represented as mean ± SEM. D. Description of 3D-Unet approach for detecting cell centroids (CC3D: 3D Connected Component Analysis). E-F. Comparison of cell detection precision (E) or recall (F) at the indicated *xy* resolutions (μm/pixel). Examples of misclassification instances contributing to false positive errors (E) or false negative errors (F) are shown above. Data represented as mean ± standard deviation. G. Cell centroids of WT cortical nuclei predicted by 3D-Unet.

We processed one brain hemisphere from four wild-type (WT) and four *Top1* cKO mice at P15 -when the *Top1* cKO had displayed large, bilateral deficits in brain structure (Fragola et al., 2020). We labeled layer-specific cell-types using antibodies for Cux1 (upper layer neuron marker) and Ctip2 (lower layer neuron marker) in addition to staining all cell nuclei with TO-PRO-3 (TP3) during iDISCO+ processing. After clearing, samples were imaged using the Ultramicroscope II -one of the most widely used commercial light-sheet microscopes for imaging cleared tissues (Cai et al., 2019; Ertürk et al., 2012; Kirst et al., 2020; Liebmann et al., 2016; Pan et al., 2016; Renier et al., 2016; Susaki et al., 2015; Tainaka et al., 2014; Ye et al., 2016). The *Top1* cKO hemispheres displayed a noticeable reduction in overall cortical volume (Figure 1B). During light-sheet imaging, there is a well known trade off between optical resolution, particularly in the axial (*z*) dimension, and imaging speed. While the Ultramicroscope II features axial sweeping to maintain relatively even *z* resolution throughout the field of view (Dean et al., 2015), the additional mechanical movement of the light-sheet significantly reduces the imaging rate. After testing various imaging schemes, we imaged at 1.21×1.21×4 (μm/voxel) resolution with a light-sheet thickness of 9 μm. The resulting images provided sufficient resolution to visually delineate cell nuclei in the cortex (Figure 1C) while limiting imaging time to 10-15 hours for all 3 channels in 1 WT hemisphere (∼9 hours for *Top1* cKO).

Prolonged imaging of cleared tissue samples can induce several artifacts over the course of image acquisition. In particular, drift in the sample or aberrant microscope stage movement can cause misalignment between image tile positions within and between channels. These issues become more pronounced at higher optical resolution where slight variations can prevent colocalization of cell nuclei with their respective immunolabeled markers. To ensure correct alignment between channels, we applied a series of rigid and non-rigid registration steps using the Elastix toolbox (Klein et al., 2010) to map the Cux1 and Ctip2 channels onto the TO-PRO-3 channel without inducing non-specific local background warping (Figure S1). We also found that many of the commonly used programs for performing 3D image stitching (Bria and Iannello, 2012; Hörl et al., 2019) did not accurately align adjacent tile stacks due to spurious stage movement, which has been noted by other groups (Kirst et al., 2020). To ensure accurate image reconstruction, we applied a simplified iterative 2D stitching procedure that uses scale-invariant feature transforms (Lowe, 2004) to produce continuous images without cell duplication along tile edges (Figure S2). Finally, differences in fluorescence intensity caused by light attenuation and photo bleaching during the course of imaging can result in uneven brightness between image tile positions. To ensure uniform signal across tiles, we measured the differences in image contrast in overlapping tile regions to estimate and correct for variations in signal intensity among tile stacks (Figure S1D).

Completion of the preprocessing steps described above resulted in aligned, fully stitched 3 channel images and datasets <1TB per sample (∼400GB for WT and ∼180GB for *Top1* cKO). The *Top1* cKO hemispheres displayed clear reductions in thickness throughout the cortex (Figure 1D). While all cortical layers showed some amount of degeneration, Layer 5 and Layer 6 neurons seemed to be more severely depleted (Figure 1E) and we hypothesized that certain cortical areas may be differentially impacted as well.

### Point Correspondence Improves Image Registration for Structures with Large Morphological Differences

Because of the significant differences in gross morphology within the *Top1* cKO brain, image registration was not accurate using only intensity-based mutual information metrics (Figure 2A). To improve registration accuracy of the *Top1* cKO brain, we manually selected up to 200 points at distinguishable structure landmarks in the Nissl stained ARA and their corresponding locations in the TO-PRO-3 nuclei channel for each sample. Point locations were positioned primarily around the cortex as this was our region of interest (Figure S3A). Using Euclidean point distances as an additional metric during the registration process significantly improved cortical annotation when compared to a manually delineated mask (Figure 2A). Increasing the number of points resulted in higher DICE similarity coefficient scores in *Top1* cKO samples (*Top1* cKO MMI, mean = 0.526, s.d. = 0.189; *Top1* cKO MMI + 200 Pts, mean = 0.890 s.d. = 0.013) indicating improvements in registration accuracy (Figures 2B and S3B-D). These results show that point correspondence can be used to better register mouse models with large structural variation.

Using the spatial deformation fields generated after image registration, we analyzed which areas in the *Top1* cKO cortex exhibited the largest changes in volume relative to WT. While the cortex as a whole showed a large reduction in volume (mean = 80%, s.d. = 3.7%, p < 0.001), we observed slightly greater decreases in frontal regions, such as the orbitofrontal (ORB) and infralimibic (ILA) areas, as well as certain lateral regions near the temporal association area (TEa) (Figures 2B and 2C). This suggests that the neuronal cell-types within these structures may be more susceptible to degeneration upon *Top1* deletion.

### 3D-Unet Accurately Quantifies Cell Nuclei in the Cortex

3D cell segmentation of tissue cleared images can be difficult due to the density of cells in the brain, limits of imaging resolution, and overall data complexity. Here we implemented a deep learning model, based on a 3D version of the popular U-Net framework (3D-Unet) (Çiçek et al., 2016; Isensee et al., 2018), to accurately quantify the total number of cell nuclei marked by TO-PRO-3 staining within the cortex. We generated two sets of manually labeled nuclei: (1) For training, ∼67,000 cortical nuclei were manually delineated from 256 training image patches (112×112×32 voxels/patch) of cortical nuclei at either high (0.75×0.75×2.5 μm/voxel) or low (1.21×1.21×4 μm/voxel) spatial resolutions. To increase manual delineation efficiency, we focused only on cell detection by delineating a 2D binary mask at the middle Z position to be used as a marker for each cell nucleus. (2) For evaluation, an independent set of ∼3,500 manually delineated nuclei were used where the full 3D extent of the nucleus was labeled in order to determine accuracy of predicted centroid placement. Cell marker predictions within each 3D patch were then thresholded and analyzed for connected components to calculate final cell centroid positions (Figure 2D).

To evaluate cell detection accuracy, we compared precision and recall rates for detecting nuclei in the evaluation dataset using 3D-Unet and two previously published analysis tools for tissue cleared images with cell counting components: ClearMap and CUBIC Informatics (CUBIC). In our tests, 3D-Unet achieved the highest precision and recall rates in both high and low resolution images when the full training datasets were used (Figures 2E and 2F). At low resolution, 3D-Unet achieved significantly lower error rates compared to the next best performing method (CUBIC) at higher resolution (p = 0.043, CUBIC 0.75/3D-Unet 1.21; p < 0.001, CUBIC 1.21/3D-Unet 1.21; p < 0.001, ClearMap 1.21/3D-Unet 1.21; McNemar’s test).

Using the trained 3D-Unet model, we counted 8.43(± 0.05)×10^6^ cells in the P15 WT cortex (Figure 2G), which was similar to previously published results in adult mice (Murakami et al., 2018). This indicates that, with sufficient training, deep neural networks can compensate for a lack of imaging resolution and achieve accurate cell quantification.

### Lower Layer Neurons in the Frontal Cortex Are Preferentially Targeted by *Top1* Deletion

To quantify neuronal cell-types in WT and *Top1* cKO cortexes, we developed a supervised Support Vector Machine (SVM) model to classify cell-types based on local intensity, shape, and annotation features. We found that a supervised approach, after training on 1,000 nuclei in each brain sample, achieved more accurate classification compared to an unsupervised mixture model approach (Figure S5). After removing outliers and summing across cortical structures, we counted 1.74(± 0.07)×10^6^ Ctip2+ and 1.94(± 0.05)×10^6^ Cux1+ in WT compared to 0.30(± 0.08)×10^6^ Ctip2+ and 0.73(± 0.11)×10^6^ Cux1+ in the *Top1* cKO (Figures 3A and 3B). Overall, this constitutes an ∼83% decrease in Ctip2+ cells and ∼62% decrease in Cux1+ cells. When compared to previous results in 2D sections from somatosensory cortex (Fragola et al., 2020), we saw a similar bias towards lower layer neuron degeneration (Cux1/Ctip2 = 1.97 in 3D SSp; 2.33 in 2D), however with a larger reduction in total neuron counts. While this can be partially attributed to differences in cell quantification methods, the increase in sampling depth from volumetric analyses can also uncover larger effects in total cell count compared to serial 2D analysis.

**Figure 3.**
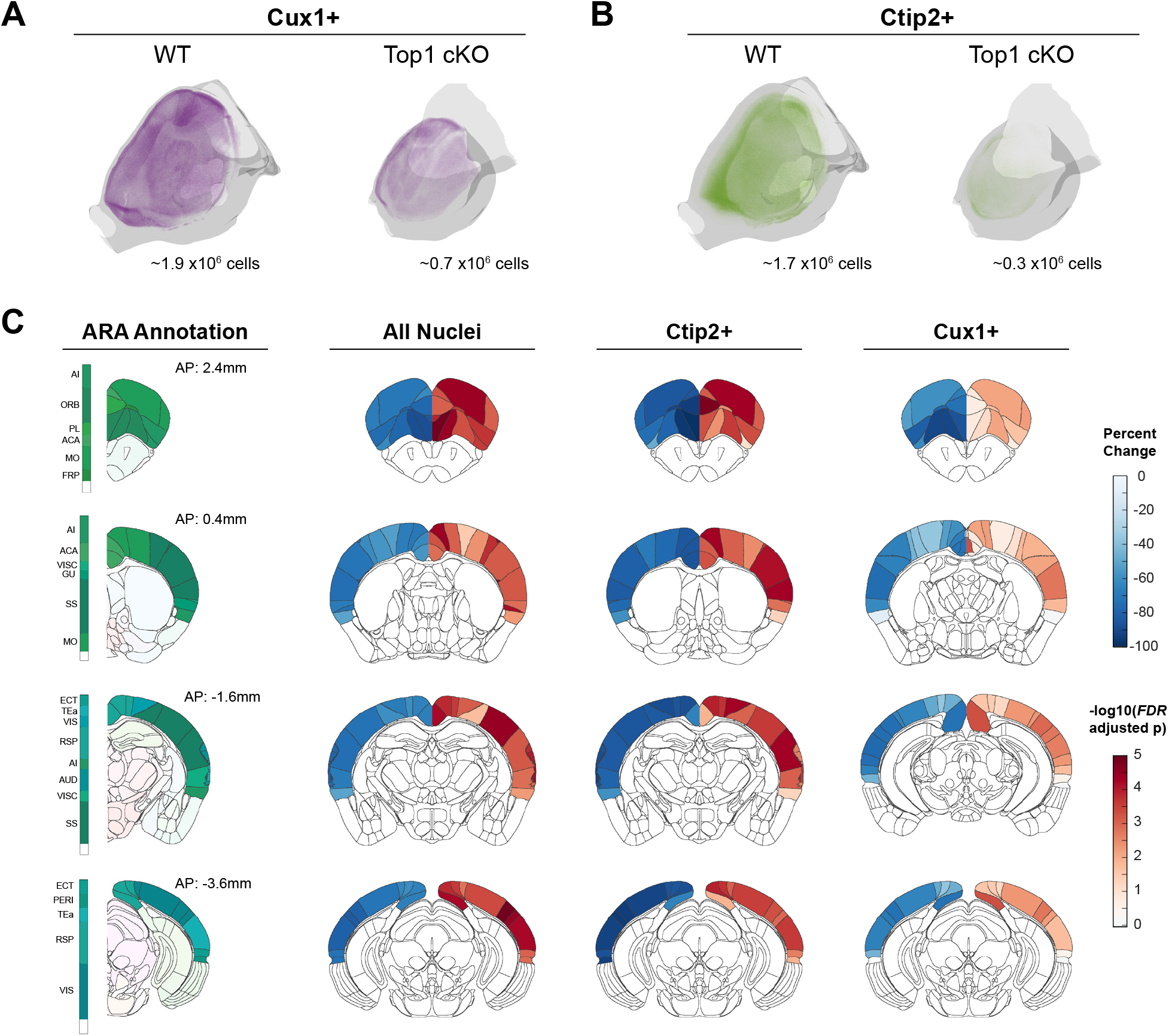
*Top1* Deletion Induces Broad Degeneration of Neuronal Cell-types Particularly in Frontal Regions. A-B. Point cloud display of Cux1+ (A) or Ctip2+ (B) cells within WT and *Top1* cKO cortexes. C. Coronal slice visualizations displaying percent change in cell count (left hemisphere) and FDR-adjusted p values (right hemisphere). Colored ARA annotations at corresponding positions displayed for reference.

Next, we compared differences in cell counts and density for 43 cortical areas defined by functional connectivity in the ARA (Harris et al., 2019) and the complete isocortex to see which regions were most affected by *Top1* deletion. After correcting for multiple comparisons (FDR < 0.05), all but one of the 43 structures showed a significant decrease in total TO-PRO-3 cell count indicating broad degeneration across all cortical areas in the *Top1* cKO model (Figures 3C and S6E). Among neuronal cell-types, we identified 25 and 41 structures with significant decreases in Cux1+ and Ctip2+ cell counts, respectively. While many structures, including several areas in somatosensory cortex (SSp-n, SS-m, SSp-bfd), shared significant losses in both Cux1+ and Ctip2+ excitatory neurons, the largest reductions were seen in Ctip2+ cells localized in frontal areas, such as the prelimbic area (PL) and secondary motor area (MOs) not measured in previous work (Fragola et al., 2020). We then calculated cell density by normalizing counts to registered structure volumes. Interestingly, the majority of structures show significant increases in TO-PRO-3+ cell density (Figure S6E), suggesting that, in addition to cell loss, degeneration of neuronal processes is also contributing to differences in cortical structure.

Structures with the largest increases were again localized in frontal regions, such as the prelimbic (PL), infralimbic (ILA), and orbitofrontal (ORB) areas, as well as medial regions, such as the anterior cingulate areas (ACA). Decreases in cell number also resulted in greater reductions in cortical surface area compared to cortical thickness (Figure S6B-D). Taken together, these results show that, even in cases where genetic perturbation induces strong phenotypic effects such as in the *Top1* cKO model, NuMorph can reveal more localized differences in cell-type number within specific brain regions.

### Neurodegeneration is Spatially Correlated with Genes Differentially Expressed in *Top1* cKO

Previous evidence suggests that lower layer neurons, particularly those in L5, are most susceptible to degeneration as a result of reduced expression of long, neuronal genes in the *Top1* cKO model (Fragola et al., 2020). While the severe structural deficits in *Top1* cKO precluded us from accurately quantifying L5 neurons in individual cortical regions, we found that regions with large L5 volumes in the ARA saw the greatest reductions in total structure volume in *Top1* cKO (Figure 4A). Furthermore, these regions also saw the largest increases in cell density (Figure 4B) suggesting local degeneration of neuronal processes. We then performed spatial correlations between regional cell count differences and gene expression using in-situ hybridization (ISH) data from Allen Mouse Brain Atlas (AMBA) (Lein et al., 2007). We tested whether the degree of *Top1* cKO induced structural change among cortical regions was related to the expression of long genes (i.e. genes >100kb) within those regions, as *Top1* is known to be a transcriptional regulator of long genes (King et al., 2013; Mabb et al., 2016). We found that in WT, regions with higher densities of Ctip2+ lower layer neurons were significantly associated with increased long gene expression (Figure 4C), providing further support that lower layer neurons express longer genes. Additionally, regions with larger reductions in cell numbers in *Top1* cKO were correlated with increased long gene expression (Figure 4D). Interestingly, fold change in Ctip2+ count differences saw the lowest positive correlation, likely because significant lower layer degeneration had already occurred by P15, minimizing variation between individual cortical regions. Gene Ontology analysis using random-null ensembles to overcome gene-enrichment bias (Fulcher et al., 2020), identified 113 functional annotations associated with greater neuronal loss, including several processes involved in axon guidance and extension (Figure S6F). We then searched for spatial correlations with individual genes differentially expressed in the P7 *Top1* cKO cortex as measured by scRNA-seq (Fragola et al., 2020). Among the 125 differentially expressed genes in *Top1* cKO that also contained ISH signatures in the AMBA, 5 were significantly correlated with relative difference in excitatory neuron count (Figure 4E). The most signficant gene, S100a10 (also known as p11), is predominantly expressed by L5a corticospinal motor neurons in the cortex (Arlotta et al., 2005; Milosevic et al., 2017). Large reductions in Ctip2+ neurons in the *Top1* cKO secondary motor area (MOs) and other frontal areas where S100a10 is highly expressed, suggest that changes in S100a10 expression may increase susceptibility for L5 degeneration in these regions (Figures 4F and 4G). These results demonstrate how existing spatial gene expression resources can be leveraged with cleared tissue analysis to identify the specific genes, cell-types, and biological processes contributing to gene-structure associations.

**Figure 4.**
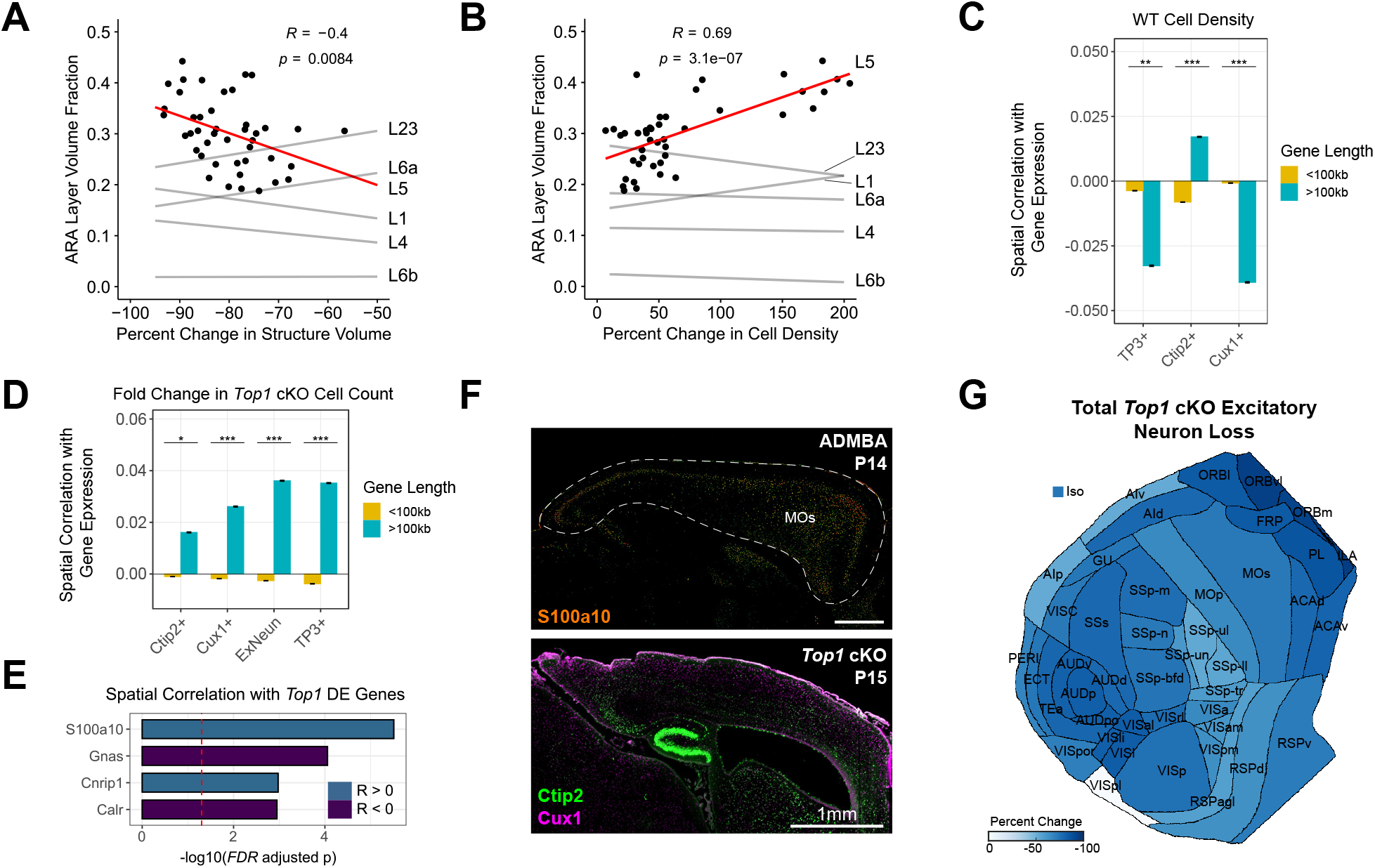
Effects of *Top1* Deletion Are Associated with Spatial Patterns of Gene Expression. A-B. Association between structure volume (A) and cell density (B) differences in *Top1* cKO with L5 volume as a fraction of total volume in the ARA. Data points shown for L5 associations. (R: Pearson correlation coefficient). C-D. Association between spatial gene expression and WT cell density (C) or negative fold change in cell count between *Top1* cKO and WT (D). Spearman correlation coefficients, binned by gene length, for each gene’s expression across cortical regions were used for comparisons. Increased correlation indicates stronger association with cell loss in (D). TP3+ indicates all cells and ExNeun indicates excitatory neurons (i.e. Ctip2+ or Cux1+). Displaying mean ± SEM. E. Genes differentially expressed in *Top1* cKO excitatory neurons significantly correlated with relative change in excitatory neuron count across cortical regions (Spearman; FDR < 0.05). F. ISH expression of S100a10 at P14 in the Allen Developing Mouse Brain Atlas (ADMBA) with the cortex outlined and a corresponding sagittal section of *Top1* cKO. (MOs: secondary motor area). G. Flattened isocortex displaying percent change in excitatory neuron counts (i.e. Ctip2+ or Cux1+) in *Top1* cKO relative to WT.

### Conditional *Nf1* Deletion Induces a Brain Overgrowth Phenotype that Shares Similarities with Human MRI Results

While the *Top1* cKO model served as a suitable test case for applying NuMorph to study severe brain structure deficits, germline loss-of-function mutations in *Top1* are highly deleterious and are extremely rare in humans (Karczewski et al., 2020). To further validate the utility of NuMorph for analyzing more subtle structural phenotypes in a disease-relevant animal model, we applied NuMorph to investigate *Nf1* knockout models of Neurofibromatosis type I (NF1). We generated two novel *Nf1* conditional knockout mouse models with one (*Nf1*^*fl/+*^;*Emx1-Cre*) or both (*Nf1*^*fl/fl*^;*Emx1-Cre*) copies of the *Nf1* gene conditionally deleted in dorsal telencephalic progenitor cells by *Emx1* promoter-driven Cre-recombination (Gorski et al., 2002). We chose *Emx1:Cre* mouse line because Cre recombinase activity can be detected as early as E10.5, 1 and 2 days earlier than Nestin:Cre (Tronche et al., 1999) and hGFAP-Cre lines (Zhuo et al., 2001), respectively. In addition, the expression of Cre recombinase in *Emx1:Cre* mice is highly restricted to the dorsal telencephalic progenitor cells that allows us to investigate the cortex-specific effect of *Nf1* deletion. We found that bi-allelic *Nf1* inactivation resulted in increased brain weight and decreased body weight compared to control and mono-allelic inactivation (Fig. S7A-C). The increase in brain weight is evident as early as P0, suggesting a possible alteration in cortical development at the embryonic stage, a time window which is critical for both cortical neurogenesis and gliogenesis. We sought to systematically characterize this brain overgrowth phenotype using NuMorph.

We performed tissue clearing and whole brain imaging of six control, six heterozygous knockout (*Nf1*^*fl/+*^;*Emx1-Cre*), and six homozygous knockout (*Nf1*^*fl/fl*^;*Emx1-Cre*) brain hemispheres in littermate groups using the same Ctip2/Cux1 antibody panel as in the *Top1* study. *Nf1*^*fl/fl*^;*Emx1-Cre* showed typical localization of Ctip2+ lower layer neurons and Cux1+ upper layer neurons (Fig. 5A,B). Using NuMorph we also measured the average cortical thickness of 17 cortical regions and detected increased thickness in posterior regions such the visual (VIS), auditory (AUD), and posterior parietal association areas (PTLp) with slight cortical thinning in orbitofrontal (ORB) areas in the *Nf1*^*fl/fl*^;*Emx1-Cre* model (Fig. 5C). These differences bear a strikingly similar pattern to human MRI findings where patients with neurofibromatosis type 1 were shown to have thicker occipital and thinner frontal cortices (Barkovich et al., 2018). Much of the increase in overall cortical thickness was driven by expansion of cortical layers 5 and 6 (Fig. S7D). To identify which cell-types were leading to increased cortical thickness in these regions, we quantified the number of ToPro3+ nuclei throughout the cortex and found a noticeable increase in overall cell count in *Nf1*^*fl/fl*^;*Emx1-Cre* (25% increase, p = 0.011) that was largely attributed to greater numbers of Ctip2-/Cux1-non-excitatory neuron cell-types (50% increase, p = 0.001) (Fig. 5D). No significant differences in global cortical cell count were observed in the heterozygous *Nf1*^*fl/+*^;*Emx1-Cre* model.

**Figure 5.**
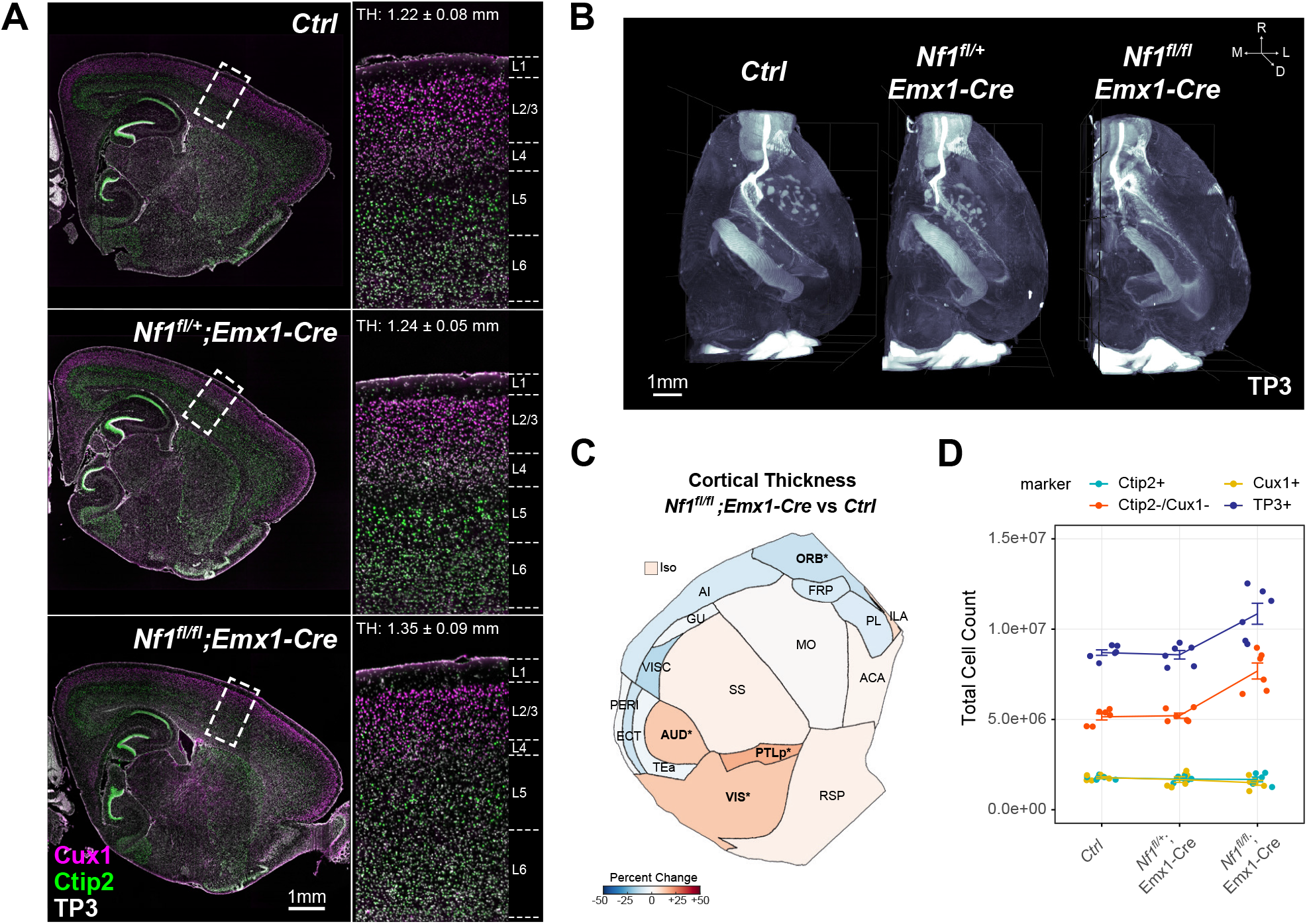
*Nf1* Deletion Induces Cortical Thickening Driven by Increased Numbers of Non-Excitatory Neuronal Cell-types. A. Optical sagittal sections of immunolabeled lower layer (Ctip2+) and upper layer (Cux1+) neurons in P14 *Ctrl, Nf1*^*fl/+*^;*Emx1-Cre*, and *Nf1*^*fl/fl*^;*Emx1-Cre* brain hemispheres. Zoomed in regions of boxed cortical areas near somatosensory cortex showing expected localization of upper and lower layer neurons. Average cortical thickness (TH) measurements indicated for full 3D somatosensory volumes. B. 3D rendering of cell nuclei in *Ctrl, Nf1*^*fl/+*^;*Emx1-Cre*, and *Nf1*^*fl/fl*^;*Emx1-Cre* brain hemispheres. C. Flattened isocortex displaying percent change in cortical thickness in *Nf1*^*fl/fl*^;*Emx1-Cre* across 17 broad regions and the full isocortex (Iso) compared to *Ctrl*. Significant regions are bolded (FDR<0.05) and starred. Structure name abbreviations provided in Table S1. D. Total isocortex counts of each cell-type class measured across all *Nf1*^*+/+*^, *Nf1*^*fl/+*^;*Emx1-Cre*, and *Nf1*^*fl/fl*^;*Emx1-Cre* samples.

### Astrocytes and Oligodendrocytes Drive Increased Cortical Thickness in the *Nf1*^*fl/fl*^*;Emx1-Cre* Model

We further investigated which specific cell-types comprised the Ctip2-/Cux1-class of cells that was driving cortical expansion in the *Nf1*^*fl/fl*^;*Emx1-Cre* model and how these effects varied across cortical regions. We found broad increases in the proportion of Ctip2-/Cux1-cells throughout the cortex (40/44 structures FDR<0.05) with the greatest increases seen in posterior and medial areas such as the Retrosplenial, Auditory, and Visual cortices (Fig. 6A). As expected, regions with higher Ctip2-/Cux1-cell numbers showed a significant positive correlation with cortical thickness (Fig. S7E). To identify the specific cell-types within the Ctip2-/Cux1-class, we focused on non-neuronal cell-types differentiated from the *Emx1-Cre* expressing lineage: astrocytes and oligodendrocytes. Based on previously estimated cell type proportions within the adult mouse brain (Erö et al., 2018), regions with larger increases in Ctip2-/Cux1-cell numbers in the *Nf1*^*fl/fl*^;*Emx1-Cre* model typically have a higher fraction of glial cell-types in the wild-type cortex (Fig. 6B), suggesting that the non-excitatory neuronal cells were glia. Furthemore, previous studies have shown inactivation of *Nf1* results in aberrant proliferation of both astrocytes and oligodendrocytes (Bajenaru et al., 2001; Gutmann et al., 1999; Hegedus et al., 2007; Wang et al., 2012). To confirm an expansion in glial cell numbers was present, we performed immunolabeling of P14 sections taken from the somatosensory cortex for Olig2 and GFAP, markers of oligodendrocytes and astrocytes, respectively, where we detected increased numbers of both Olig2+ and GFAP+ cells in this region (Fig. 6C-F). Combined with our measurements of cortical thickness, these results suggest that increased production of glial cell-types may explain the cellular mechanism that leads to increased thickness of posterior cortical regions observed in NF1 patients.

**Figure 6.**
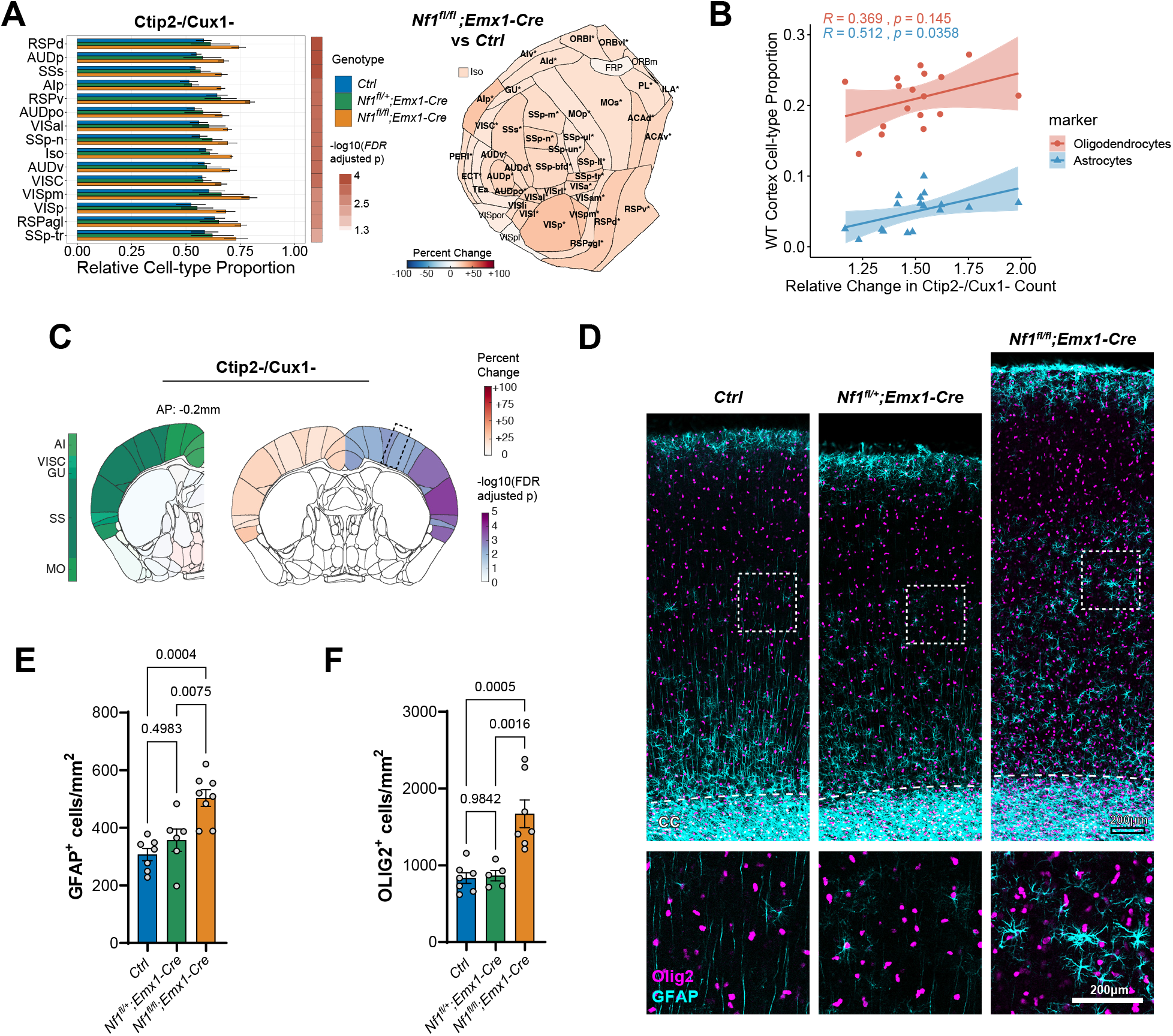
*Nf1* Deletion Increases Production of Astrocytes and Oligodendrocytes Broadly Throughout the Cortex. A. Differences in relative proportion of non-excitatory neuronal cell-types (i.e. (Ctip2-& Cux1-)/ToPro3+) across 43 cortical regions and the full isocortex (Iso) after *Nf1* deletion. The top 15 regions sorted by binned p-value and fold change are shown (*Nf1*^*fl/fl*^;*Emx1-Cre* vs. *Ctrl*, FDR<0.05). Flattened isocortex displaying percent change in *Nf1*^*fl/fl*^;*Emx1-Cre* are shown on the right. Significant regions are bolded and starred (FDR<0.05). B. Association between estimated astrocyte and oligodendrocytes proportion across regions in the wild-type cortex (Erö et al., 2018) and relative change in non-excitatory neuronal cell-types measured in *Nf1*^*fl/fl*^;*Emx1-Cre*. (R: Pearson correlation coefficient). C. Coronal slice visualization displaying percent change in cell count (left hemisphere) and FDR-adjusted p values (right hemisphere) as measured by 3D analysis.

### *Nf1* Deletion Results in a Regionally-Specific Imbalance of Upper Layer and Lower Layer Neurons

Finally, we looked at whether the proportion of Ctip2+ and/or Cux1+ excitatory neurons was altered across cortical regions following *Nf1* deletion. While only a small number of regions were significant for changes in raw cell count for either Ctip2 or Cux1 in *Nf1*^*fl/fl*^;*Emx1-Cre* compared to control, we detected stronger effects when normalizing individual counts by total cell number or by volume (Fig. 7A,B and Fig. S7G). In other words, alterations in cell-type distribution become more apparent when accounting for the increased rate of gliogenesis in the *Nf1*^*fl/fl*^;*Emx1-Cre* model. Across the entire cortex, we observed a significant reduction in the relative number of excitatory neurons following biallelic *Nf1* deletion with a slightly greater reduction in the fractional proportion of Cux1+ neurons (35% decrease, p = 0.002, 19/44 structures FDR<0.05) compared to Ctip2+ neurons (24% decrease, p = 0.007, 30/44 structures FDR<0.05). The organization of upper layer barrel fields was also disrupted in the *Nf1*^*fl/fl*^;*Emx1-Cre* model (Fig. S7H), similar to previous work (Lush et al., 2008). In addition, we observed a strong inverse relationship in the change in Ctip2/Cux1 ratio where regions with a greater reductions in Cux1+ neurons saw lower reductions or increases in the number of Ctip2+ neurons, and vice versa. This negative correlation persists when comparing either total cell counts or cell density (Fig. 7C, and Fig. S7G). These opposing findings across the cortex show the utility of a whole brain imaging approach, because slices within specific regions may not be representative of all the effects. To further investigate whether upper layer neurogenesis is disrupted in the *Nf1*^*fl/fl*^;*Emx1-Cre* model, we performed EdU pulse labeling of E16.5 mice and quantified their differentiated progeny within P14 brain sections of somatosensory cortex (Fig. 7D). We saw a significant reduction of Satb2+ upper layer neurons co-labeled with EdU indicating decreased upper layer neurogenesis at E16.5. Considering the increase in gliogenesis seen in the *Nf1*^*fl/fl*^;*Emx1-Cre* mouse, this data suggests that *Nf1* deletion may accelerate the brain’s developmental trajectory which can result in widely disparate effects on neuronal cell-type composition across cortical regions in the adult brain.

**Figure 7.**
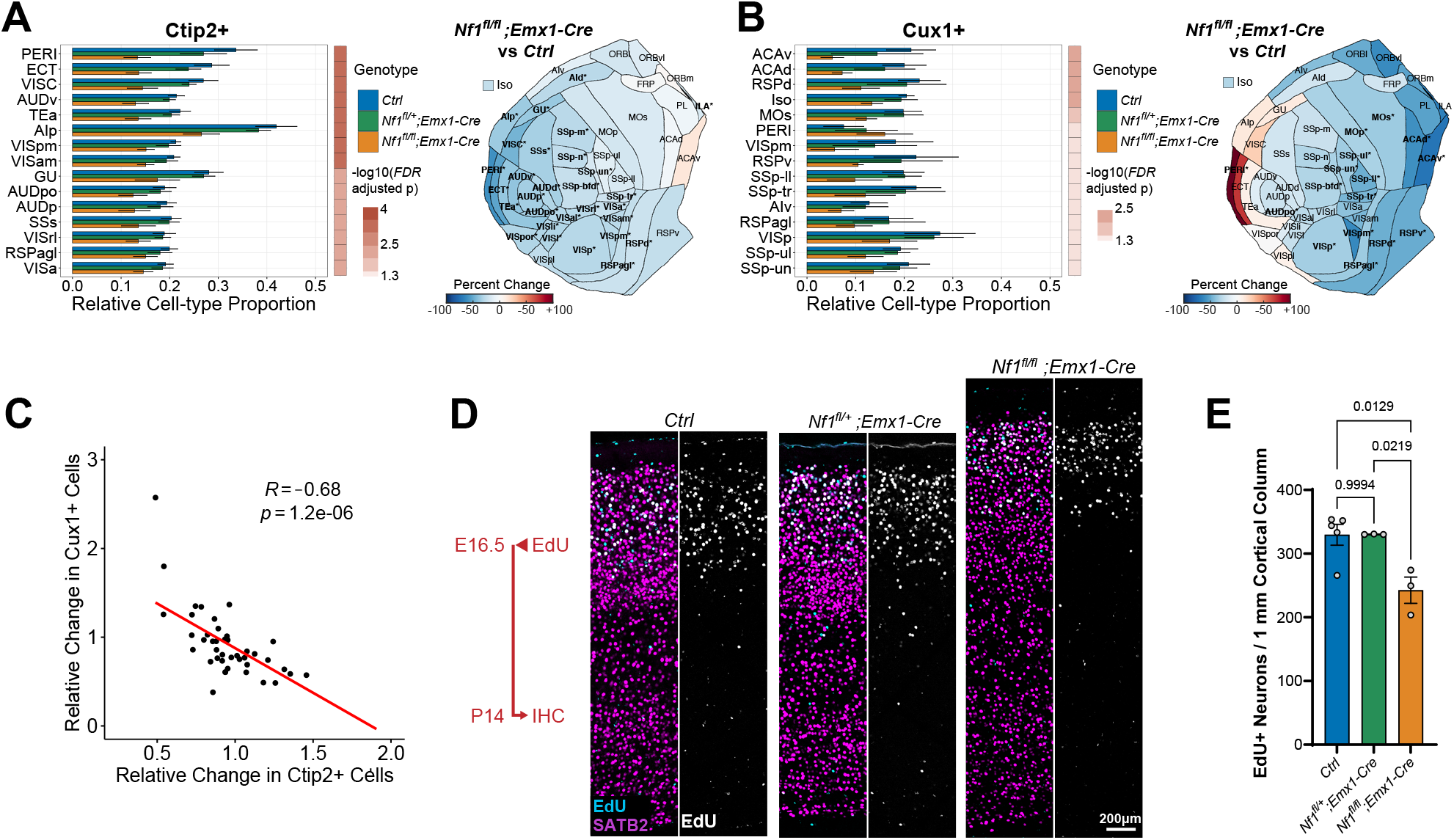
*Nf1* Deletion Alters Neuronal Cell-type Proportions in a Region-Specific Manner. A-B. Differences in relative proportion of Ctip2+ (A) and Cux1+ (B) cells across 43 cortical regions and the full isocortex after *Nf1* deletion. The top 15 structures sorted by binned p-value and fold change are shown (*Nf1*^*fl/fl*^;*Emx1-Cre* vs. *Ctrl*, FDR<0.05). Flattened isocortex displaying percent change in *Nf1*^*fl/fl*^;*Emx1-Cre* are shown on the right. Significant regions are bolded and starred (FDR<0.05). C. Association between relative change in Ctip2+ and Cux1+ cell numbers across cortical regions in the *Nf1*^*fl/fl*^;*Emx1-Cre*. D. Representative 2D sections of 1mm cortical columns from P14 somatosensory cortex after EdU injection at E16.5 showing colocalization with SATB2 (callosal projection neurons) immunolabeling in upper layers. E. Quantification of EdU+/SATB2+ cells within 1mm cortical columns in D. (n=3-5 animals, mean ± SEM).

## Discussion

Tissue clearing methods provide a unique opportunity to explore the cellular organization of the entire 3 dimensional brain structure. However, the current computational tools for analyzing cell-types in tissue cleared images have either been applied to sparse cell populations where segmentation is less difficult (Renier et al., 2016; Yun et al., 2019) or taken advantage of tissue expansion and custom-built light sheet systems to increase spatial resolution (Matsumoto et al., 2019; Murakami et al., 2018). Here, we present NuMorph, a computational pipeline for processing and quantifying nuclei within structures of the adult mouse brain acquired by conventional light-sheet fluorescence microscopy.

In the course of developing NuMorph and an appropriate imaging protocol, a large emphasis was placed on outlining a reasonable compromise between cell detection accuracy, imaging time, and computational resources. With the imaging parameters used to resolve cortical nuclei in this study, WT brain hemispheres required 3-6 hours of imaging per channel, while end-to-end processing and analysis using NuMorph required ∼1 day with a GPU-equipped workstation. By training a 3D-Unet model on a diverse set of manually labeled nuclei from multiple imaging experiments, we were able to achieve effectively equivalent error rates at this resolution compared to 1.6x higher resolution (p = 0.91, 3D-Unet 0.75/3D-Unet 1.21; McNemar’s test) that would have otherwise required significantly longer imaging times and expanded data size by ∼4x for a whole hemisphere acquisition. We expect cell detection accuracy using the training dataset generated here will remain high for analyzing other brain regions with similar cell density, while supplementation with additional training data may be needed for denser structures such as the hippocampus. Furthermore, NuMorph provides additional features and flexibility such as (1) targeting analyses to specific structures after registration to avoid unnecessary computation time, (2) detecting cells directly by nuclear protein marker expression without DNA staining, and (3) classifying cell-types by cellular markers using either supervised or unsupervised methods.

Top1 is critical for maintaining genomic stability and regulating the expression of long genes important for neuronal function (McKinnon, 2016). Recent evidence suggests that many of these same long genes contribute to neuronal diversity and have the greatest expression in the forebrain (Sugino et al., 2019). In the developing cortex, scRNA-seq studies found that L5 neurons had higher long gene expression compared to neurons from other cortical layers (Loo et al., 2019). In this study, we found that *Top1* deletion preferentially targeted many frontal areas with high L5 thickness, larger numbers of Ctip2+ lower layer neurons, and greater long gene expression. These effects likely occur much earlier than the time point studied here as previous behavioural assays showed that severe motor deficits are present as early as P7 (Fragola et al., 2020). Interestingly, inhibition of S100a10 -the gene most correlated with neuron loss -was recently shown to have a neuroprotective effect, delaying motor neuron loss in a mouse model amyotrophic lateral sclerosis (ALS) (García-Morales et al., 2019). Because *Top1* deletion results in multiple stress factors that negatively impact cell health, additional studies will be needed to disambiguate which mechanisms ultimately lead to biased degeneration of certain neuronal subtypes across brain regions.

Human MRI studies have detected gross cortical structural differences in individuals with neuropsychiatric disorders as compared to neurotypical controls (van Erp et al., 2018). The cellular basis underlying these differences cannot be assessed with standard *in vivo* MRI, due to the low resolution and lack of cellular labels. Overall increases in brain size and regional variabilities in cortical thickness were previously detected in individuals with neurofibromatosis type 1 (Barkovich et al., 2018; Payne et al., 2010). To explain the cellular basis underlying those findings, we used an approach complementary to MRI in humans, 3D cellular resolution imaging in a mouse model. Our *Nf1* knockout model exhibited a broad expansion in glial cell numbers that drove cortical thickness increases particularly in posterior regions which reproduced human MRI measurements. Numerous studies have reported increased glial cell proliferation following *Nf1* inactivation in both animal models and human iPSC lines (Gutmann et al., 2017; Hegedus et al., 2007; Wang et al., 2012; Zhu et al., 2001). However, this is the first study, to our knowledge, that performs a systematic comparison of areal differences upon *Nf1* inactivation and how these relate to changes in brain structure seen in patients with NF1. We also found striking areal differences in upper and lower layer neuron proportions upon *Nf1* biallelic deletion in cortical neural progenitor cells early in development. This highlights the key advantages of 3D whole brain imaging over 2D sectioning as stereological analysis may lead to highly variable results based on the anatomical location from which tissue is sampled. Further coupling of immediate early gene immunolabeling with cleared tissue analysis could reveal how specific structure-function relationships are altered at a cellular level in the *Nf1* and similar models (Renier et al., 2016, 2017; Ye et al., 2016).

While NuMorph has proven to be effective in analyzing moderately dense tissues such as the adult mouse cortex, the development of additional computational tools may be required to pursue more challenging experimental designs. For example, structures in the embryonic brain are typically of much higher cell density and vary in gross morphology across developmental time, making both cell quantification and image registration more difficult. In addition, segmentation and mapping of fine structures, such as neuronal processes, can be challenging with limited imaging resolution. Technological improvements in the next generation of light-sheet systems can ultimately allow for quantitative interrogation of subcellular structures at high throughput (Migliori et al., 2018; Voleti et al., 2019). However, computational tools using deep neural networks have also proven to be effective in executing diverse segmentation tasks (Friedmann et al., 2020; Kirst et al., 2020; Schubert et al., 2019; Stringer et al., 2020) or even enhancing image quality (Weigert et al., 2018). Nevertheless, community-based efforts may be needed to generate sufficient annotation data for training deep learning models to accurately perform these tasks (Borland et al., 2021; Roskams and Popović, 2016). Together we hope these new imaging and computational tools will lead to greater adoption of tissue clearing methods for quantitative analyses, rather than qualitative visualizations, of how the entire brain structure is changed by genetic or environmental risk factors for neuropsychiatric disorders.

## Acknowledgments

This work was supported by NSF (ACI-16449916 to JLS, GW, and AK), NIH (R01MH121433, R01MH118349, R01MH120125 to JLS; R01NS110791 to GW; and R56AG058663,R35ES028366 to MJZ) and DOD (W81XWH-19-1-0402 to LX), Children’s Tumor Foundation (CTF-YIA 2015-01-013 to LX), and the Foundation of Hope (to GW). We thank Pablo Ariel of the Microscopy Services Laboratory and Michelle Itano of the Neuroscience Microscopy Core for assisting in sample imaging. The Microscopy Services Laboratory, Department of Pathology and Laboratory Medicine, is supported in part by P30 CA016086 Cancer Center Core Support Grant to the UNC Lineberger Comprehensive Cancer Center. The Neuroscience Microscopy Core is supported by P30 NS045892. Research reported in this publication was also supported in part by the North Carolina Biotech Center Institutional Support Grant 2016-IDG-1016.

## Author Contributions

JLS and OK designed the study. OK performed tissue clearing, imaging, and developed NuMorph for image analysis. GF, WDS, MJZ, and LX provided fixed mouse brain tissue. BWR and LX performed additional experiments on Nf1 cKO mice. OK generated all manual annotations for training the 3D-UNet model. EHF, JM, TL, ZH, and OK performed 3D nuclei volume tracing for evaluating the 3D-Unet model. AK provided computational infrastructure to distribute data and results. GW provided guidance on image processing. JLS and LX supervised the work. OK, LX, and JLS wrote the first draft of the manuscript, and all authors provided feedback.

