## Supplemental Figures for "NuMorph: tools for cellular phenotyping in tissue cleared whole brain images"

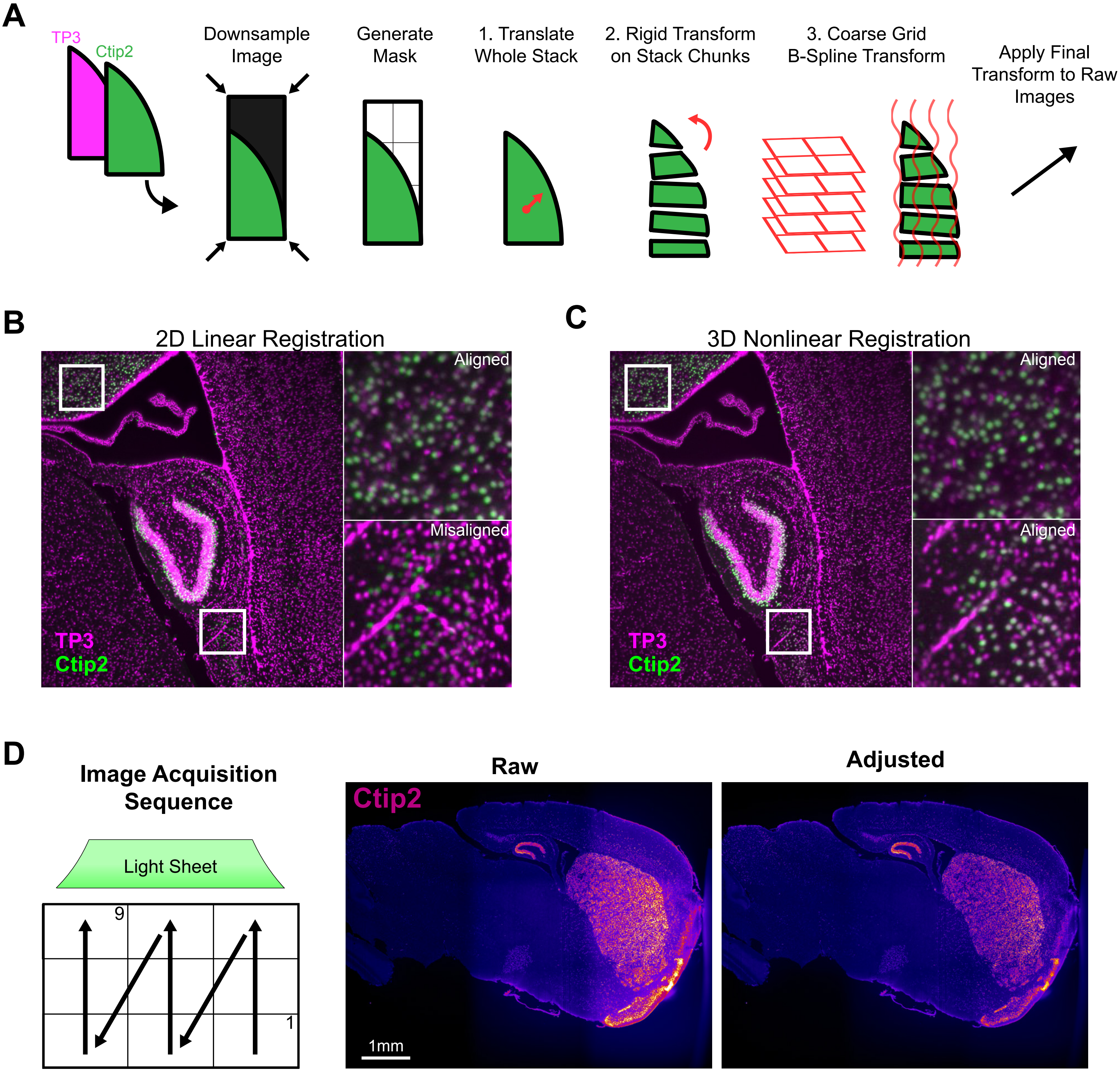


**Figure S1, Related to Figure 1. Nonlinear Alignment and Intensity Adjustment of 3D Multichannel Images.**

A. Overview of alignment procedures.

B. Example of channel misalignment after 2D registration by translation showing only part of the image aligning correctly.

C. Same section as in B after the nonlinear alignment procedure.

D. *Top1* cKO sample images of Ctip2 labeling with (right) or without (left) adjusting intensities for tile positions and light-sheet width.

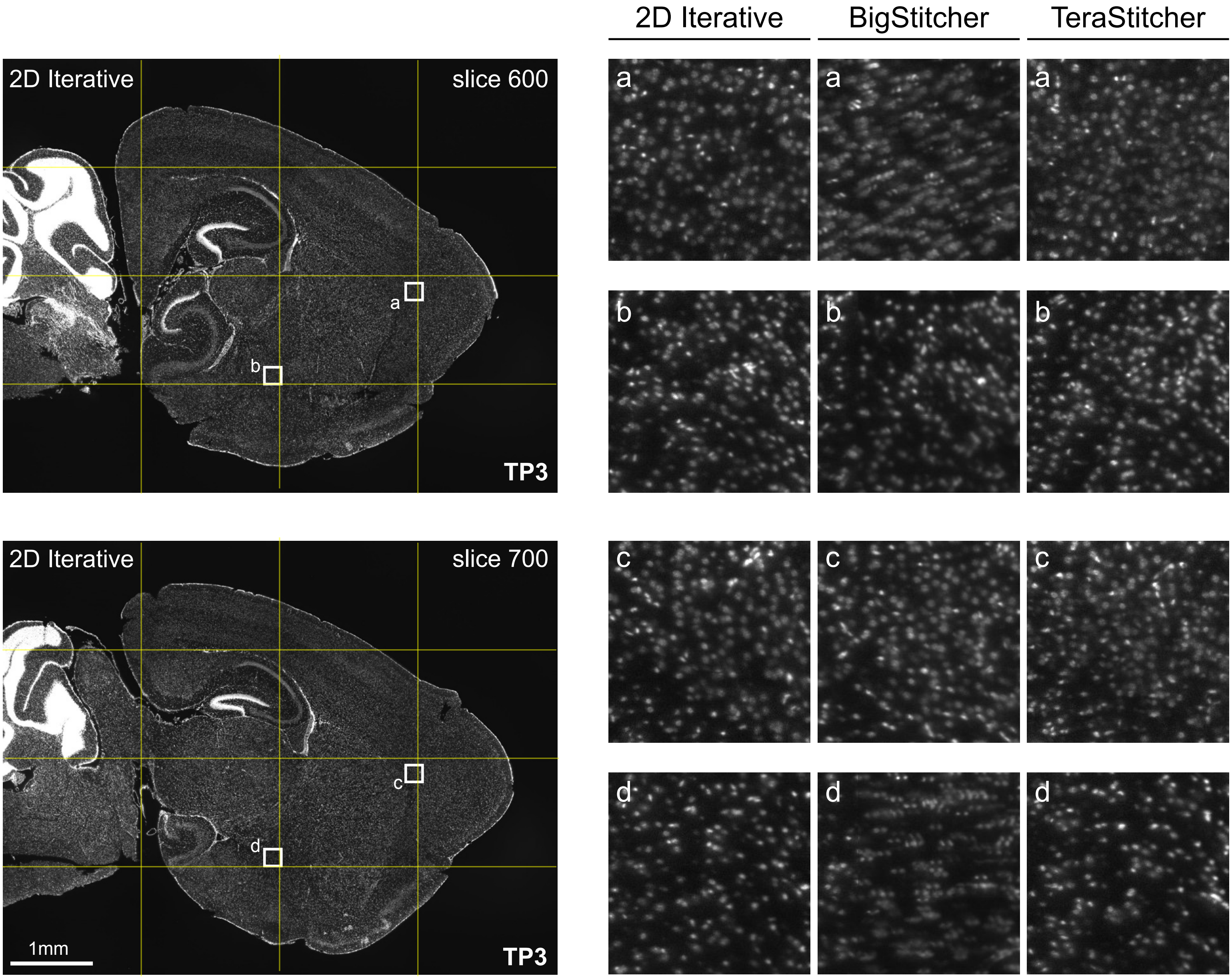


**Figure S2, Related to Figure 1. Iterative 2D Stitching of Multi-Tile Light Sheet Images.**
Sample results from 2D iterative stitching of WT mouse hemisphere compared with other dedicated 3D stitching software. Yellow lines indicate approximate stitching seams.

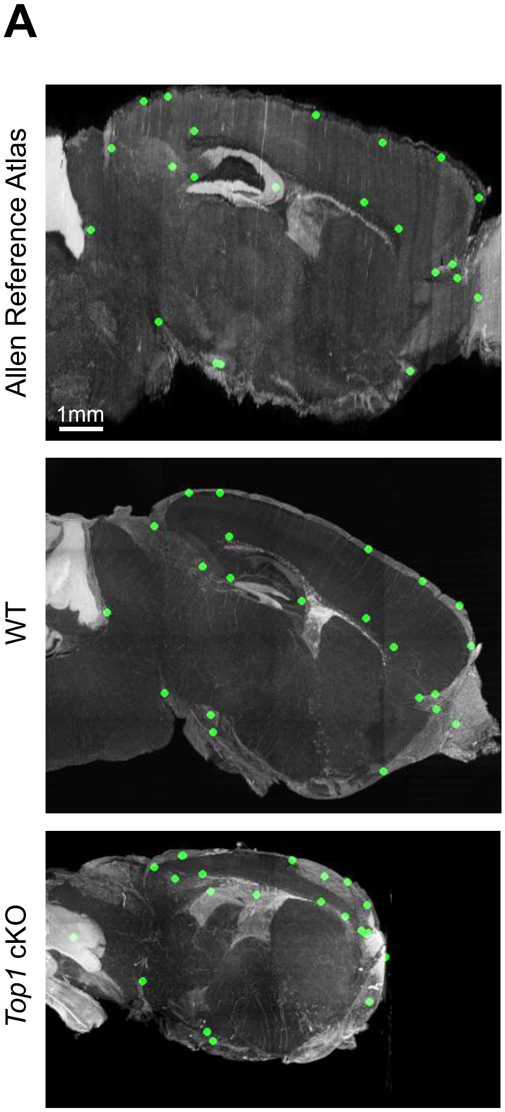


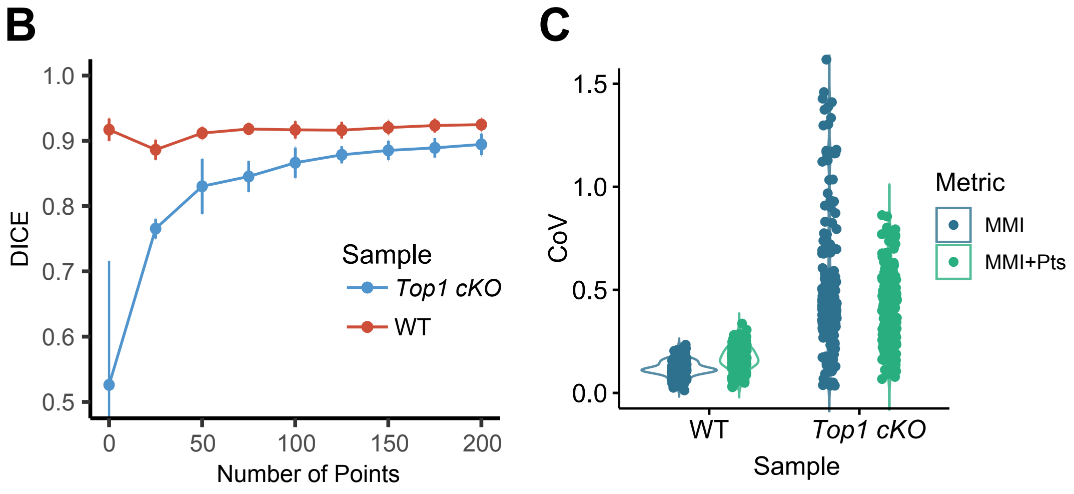


**D**

| | WT (MMI) | WT (MMI+Pts) | *Top1* (MMI) | *Top1* (MMI+Pts) |
|---|---|---|---|---|
| DICE | 0.917 (0.017) | 0.923 (0.008) | 0.526 (0.189) | **0.890 (0.013)** |
| CoV (All Cortical Structures) | 0.124 (0.037) | 0.167 (0.059) | 0.524 (0.367) | 0.422 (0.169) |
| CoV (Full Cortex Registered) | 0.096 | 0.094 | 0.203 | 0.184 |
| CoV (Full Cortex Manual) | 0.081 | | 0.242 | |

**Figure S3, Related to Figure 2. Image Landmark Selection for Points-Guided Image Registration.**
A. 1 mm thick sagittal maximum intensity projection displaying corresponding points positions in ARA, WT, and *Top1* cKO brain hemispheres.
B. DICE scores measuring cortical registration accuracy in WT and *Top1* cKO samples based on the number of points used to guide registration. Measurements with no corresponding points were made using affine + b-spline registration without a points distance metric. Data represented as mean ± standard deviation.
C. Coefficients of variation (CoV) of structure volumes for all cortical annotations in the ARA after registration with or without corresponding points.
D. DICE scores and CoV metrics for indicated registration procedures. Data represented as mean (± standard deviation). CoV was calculated for individual ARA annotations (242 structures plotted in B) or the full isocortex after registration. These compared with the CoV for the full cortex based on manual annotation. Bold value: Top1 MMI/Top1 MMI+ Pts, $p < 0.001$.

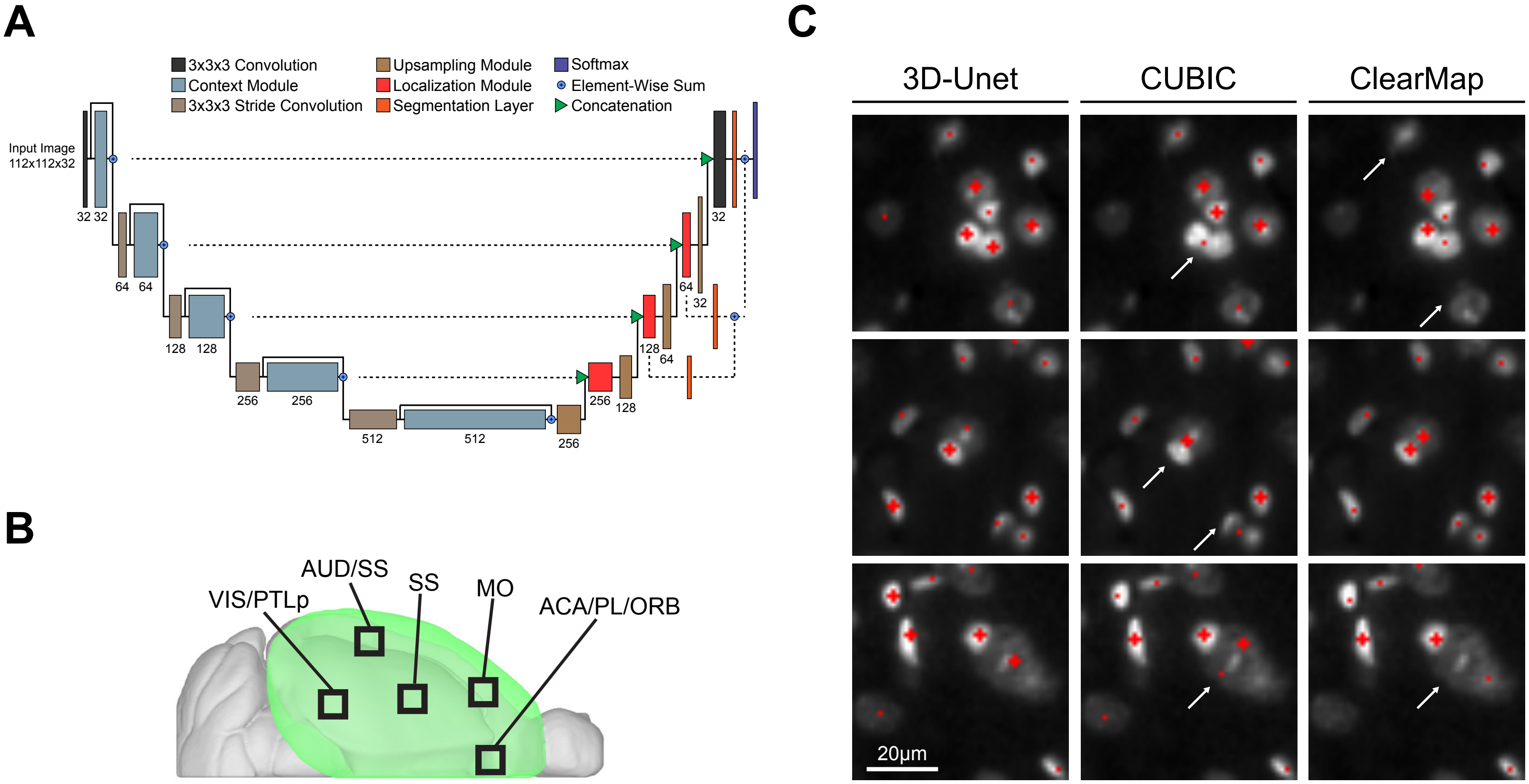


**Figure S4, Related to Figure 2. 3D-Unet Training and Evaluation.**
A. 3D-Unet architecture adapted from Isensee et al. 2018.
B. Approximate patch locations used for training the 3D-Unet nuclei detection model
C. Example images of nuclei detection results. Cross symbols indicate centroids in the displayed z slice whereas points indicate centroids in slices directly above or below. Arrows indicate detection errors in the full 3D volume.

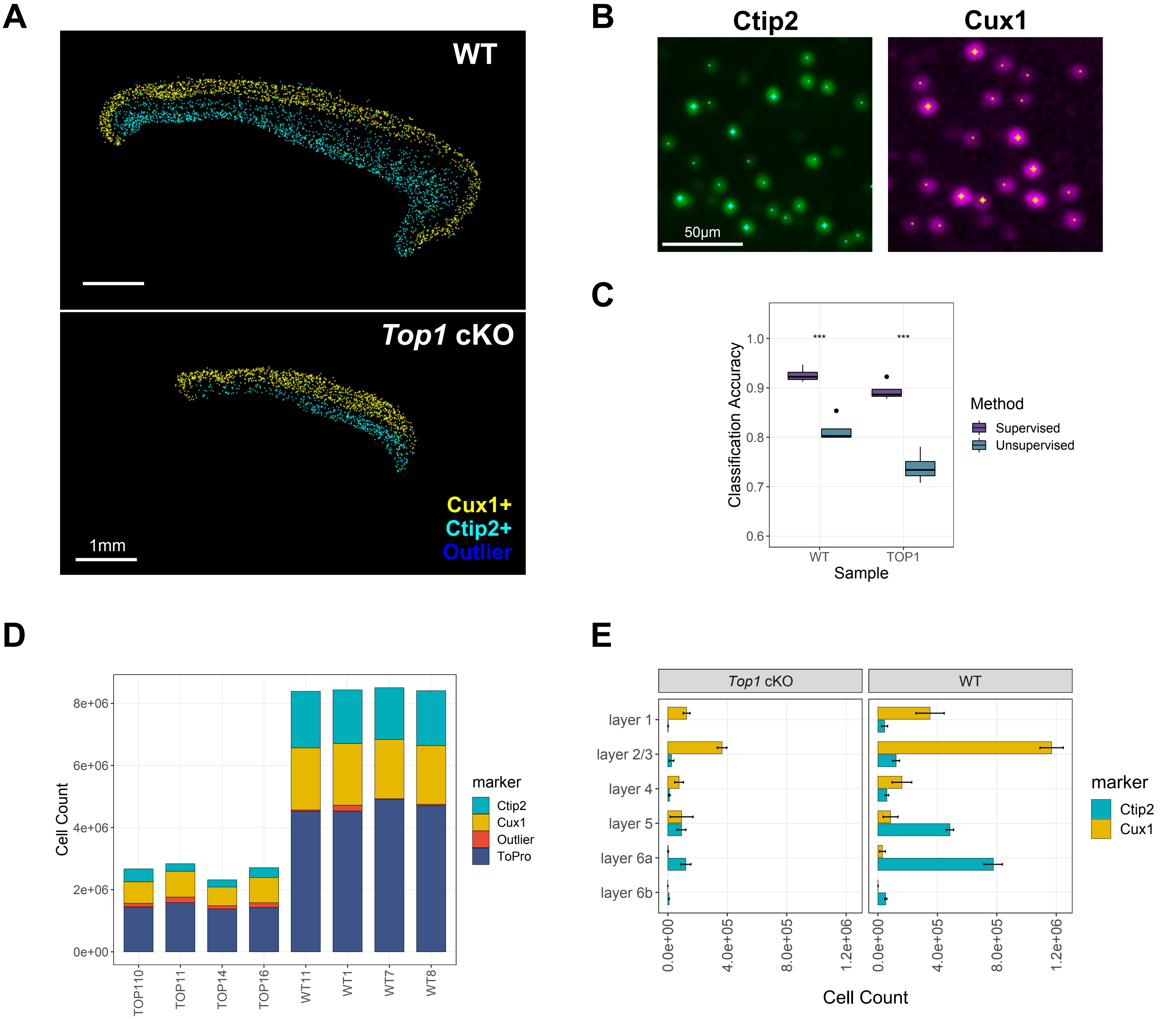


**Figure S5, Related to Figure 3. Cell-type Classification using Supervised SVM Classifier.**
A. Cell-type positions for upper and lower layer cortical neurons in a sagittal section for WT and *Top1* cKO after SVM classifications.
B. Representative images of Ctip2+ and Cux1+ cell-type classification using SVM. Cross symbols indicate centroids in the displayed z slice whereas points indicate centroids in slices directly above or below.
C. Classification accuracies using a trained SVM (supervised) classifier or by Gaussian Mixture Modeling (unsupervised). Accuracy is measured as the fraction of 1,000 cells in each sample with the correct classification based on manual identification. SVM accuracies determined based on 5-fold cross-validation. (***$p < 0.001$; McNemar test).
D. Total counts for each cell-type classification in WT and *Top1* cKO samples.
E. Distributions of Ctip2+ and Cux1+ cells across cortical layers.

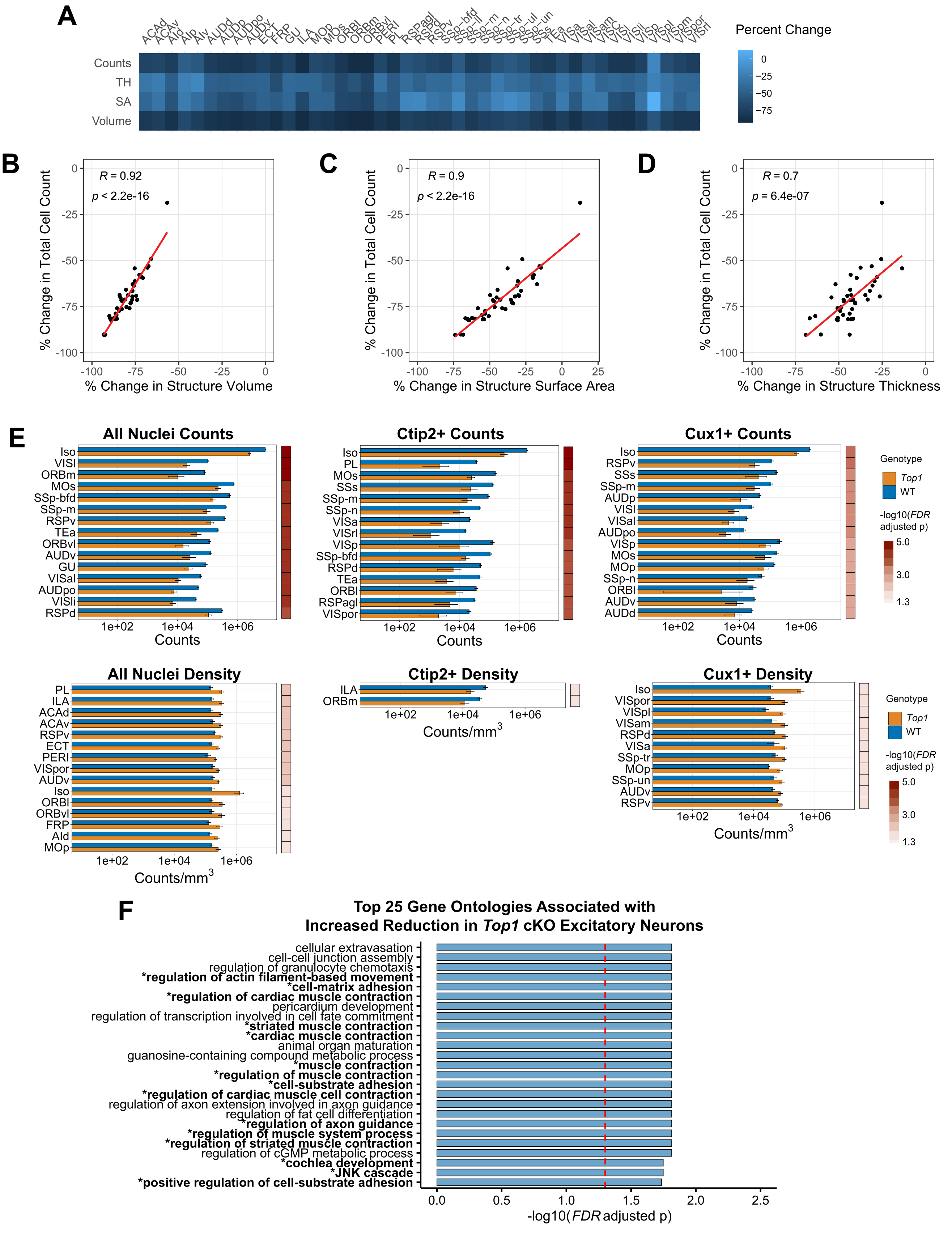


**Figure S6, Related to Figures 3 and 4. Structural and Molecular Associations with Cell Loss in the *Top1* cKO Model.**

A. Heatmap displaying percent change in cortical cell count, volume, surface area, and thickness for each cortical region.

B-D. Correlation between total cell count difference and volume (B), surface area (C), and thickness (D) across cortical regions.

E. Comparison of cell-type counts or densities between WT and *Top1* cKO across cortical regions and the full isocortex. Displaying the top 15 structures (FDR < 0.05) binned by significance level and sorted by absolute difference in count or density within each bin. Data represented as mean ± standard deviation and plotted on log10 scale. Structure name abbreviations provided in Table S1.

F. Gene ontology showing the top 25 most significant categories correlated with neuron loss in *Top1* cKO. Bolded categories contain at least 1 gene differentially expressed in *Top1* cKO from scRNA-seq studies.

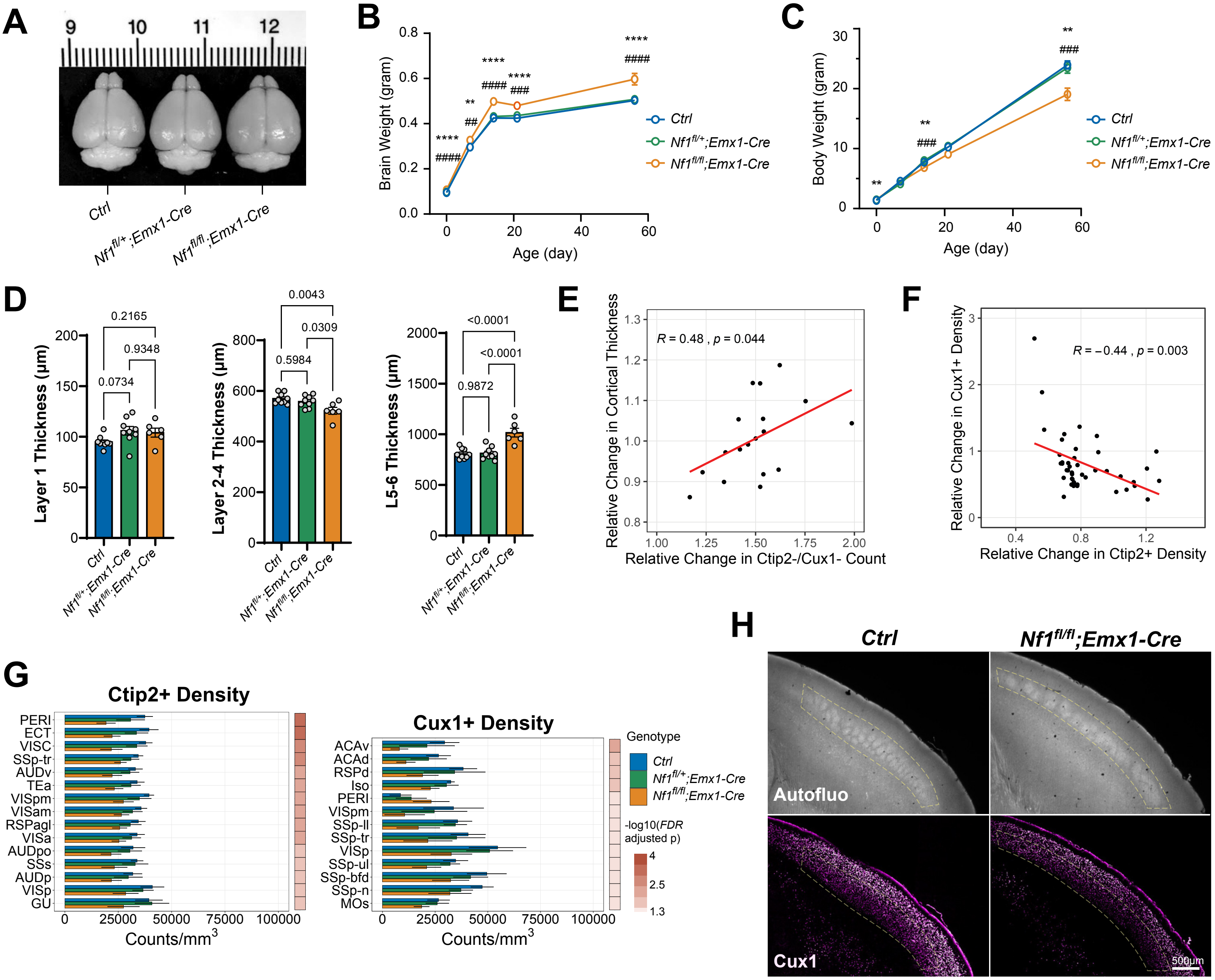


**Figure S7, Related to Figures 5-7. Characterization of *Nf1* cKO Models and Structural Associations with Cortical Cell-type Densities.**

A. Representative images of P14 brains from *Ctrl*, *Nf1$^{fl/+}$;Emx1-Cre*, and *Nf1$^{fl/fl}$;Emx1-Cre* mice.

B-C. Brain and body weight measurements for *Ctrl*, *Nf1$^{fl/+}$;Emx1-Cre*, and *Nf1$^{fl/fl}$;Emx1-Cre* mice at multiple developmental time points. (mean ± SEM).

D. Quantification of cortical layer thickness in *Nf1* models from 2D sections of somatosensory cortex.

E. Association between cortical thickness and relative change in Ctip2-/Cux1- cell counts in the *Nf1$^{fl/fl}$;Emx1-Cre* model. (R: Pearson's correlation coefficient).

F. Association between relative change in Cux1+ and Ctip2+ cell densities in the *Nf1$^{fl/fl}$;Emx1-Cre* model. (R: Pearson's correlation coefficient).

G. Differences in cell densities of Ctip2+ and Cux1+ cells across 43 cortical regions and the full isocortex after *Nf1* deletion. The top 15 structures sorted by binned p-value and fold change are shown (*Nf1$^{fl/fl}$;Emx1-Cre* vs. Ctrl, FDR<0.05).

H. Optical sections of cleared tissue autofluorescence showing disorganization of cortical barrel fields in *Nf1$^{fl/fl}$;Emx1-Cre* and reduced Cux1+ neuron density in surrounding upper layer regions.

**Table S1, Related to Figure 4-7. Volume, Cell Count, and Cell Density Statistics for *Top1* cKO and *Nf1* cKO models.**

**Video S1, Related to Figures 3 and 4. Visual Comparison of WT and *Top1* cKO Brain Hemispheres.** Representative examples of iDISCO-processed WT and *Top1* cKO samples labelling TO-PRO-3 (white), Ctip2 (green), and Cux1(magenta). Images were downsampled to 10 μm/voxel for smoother rendering. Cortical cell-type classifications displayed as point clouds (white: Ctip2-/Cux1-, teal: Ctip2+, yellow: Cux1+).

**Video S2, Related to Figure 3. 3D Inspection of NuMorph Nuclei Counting and Cell-type Classification.** Visualization of a 400 μm thick WT sagittal section at 1.21x1.21x4 μm/voxel resolution labelling TO-PRO-3 (white), Ctip2 (green), and Cux1(magenta). Cortical cell-type classifications displayed as point clouds (white: Ctip2-/Cux1-, teal: Ctip2+, yellow: Cux1+).

**Video S3, Related to Figure 5-7. Visual Comparison of *Ctrl* and *Nf1$^{fl/fl}$;Emx1-Cre* Brain Hemispheres.** Representative examples of iDISCO-processed *Ctrl* and *Nf1$^{fl/fl}$;Emx1-Cre* samples labelling TO-PRO-3 (white), Ctip2 (green), and Cux1(magenta). Major cortical areas are outlined within sliding, 500 μm thick coronal sections. Images were downsampled to 4 μm/voxel for smoother rendering.
